# Principled PCA separates signal from noise in omics count data

**DOI:** 10.1101/2025.02.03.636129

**Authors:** Jay S. Stanley, Junchen Yang, Ruiqi Li, Ofir Lindenbaum, Dmitry Kobak, Boris Landa, Yuval Kluger

## Abstract

Principal component analysis (PCA) is indispensable for processing high-throughput omics datasets, as it can extract meaningful biological variability while minimizing the influence of noise. However, the suitability of PCA is contingent on appropriate normalization and transformation of count data, and accurate selection of the number of principal components; improper choices can result in the loss of biological information or corruption of the signal due to excessive noise. Typical approaches to these challenges rely on heuristics that lack theoretical foundations. In this work, we present Biwhitened PCA (BiPCA), a theoretically grounded framework for rank estimation and data denoising across a wide range of omics modalities. BiPCA overcomes a fundamental difficulty with handling count noise in omics data by adaptively rescaling the rows and columns – a rigorous procedure that standardizes the noise variances across both dimensions. Through simulations and analysis of over 100 datasets spanning seven omics modalities, we demonstrate that BiPCA reliably recovers the data rank and enhances the biological interpretability of count data. In particular, BiPCA enhances marker gene expression, preserves cell neighborhoods, and mitigates batch effects. Our results establish BiPCA as a robust and versatile framework for high-throughput count data analysis.

## 1 Introduction

The discrete nature of biomolecules has driven the widespread use of count data in modern biology. Various experimental methods are based on counting unique entities such as RNA transcripts, open chromatin regions, or proteins in order to characterize biological phenomena and processes across different scales, from single cells to tissues, and entire organisms [1, 2, 3, 4]. High-throughput sequencing and imaging technologies are now regularly applied to diverse modalities to produce vast, high-dimensional count datasets at an unprecedented rate [5, 6, 7, 8, 9, 10]. However, while these datasets are readily collected, their analysis is often complicated by technical biases, noise, and inherent measurement variability associated with discrete counts [8, 11, 12, 13]. These obstacles necessitate robust data preprocessing and normalization methods.

Standard pipelines for preprocessing count data typically begin with quality-control filtering, proceed with modality-specific heuristic normalizations, and often conclude with principal component analysis (PCA) employed for dimensionality reduction and denoising [14, 15]. However, downstream analysis can be highly sensitive to the choice of normalization techniques and the rank used in the PCA step [16]. For example, for single-cell RNA sequencing, differences in preprocessing can strongly affect interpretations of the same experiment and may result in the omission of true biological signals or false discoveries due to noise [17, 18, 19, 20, 21]. At the same time, many preprocessing procedures often lack rigorous justifications and there has been disagreement about the most appropriate statistical models and estimation procedures [22, 23, 24, 25, 26, 27, 28, 29]. Furthermore, the rank selection for the subsequent PCA step is often not principled, instead relying on manual selection or subjective preference.

On a different front, several methods for the analysis of single-cell RNA sequencing data have been recently proposed based on random matrix theory [30, 31, 32, 33, 34]. Random matrix theory can describe the results of applying PCA to a purely random matrix, and shows that under the assumption of uniform noise, PCA eigenvalues follow the Marchenko-Pastur (MP) distribution [35]. However, these results are not directly applicable to count data. The aforementioned methods try to transform the data into standard Gaussians and then use the MP distribution to separate signal from the noise. These transformations are heavily based on heuristics and do not provide a guarantee of optimal fit and signal preservation. Furthermore, many count modalities exhibit varying dispersions, which could significantly skew the assumed normality. It is not clear how these methods can be generalized to these distributions. To address these issues, we developed an adaptive two-step pipeline for preprocessing non-negative count data, *biwhitened principal component analysis* (BiPCA). Like PCA, BiPCA assumes that without the noise, the data would be constrained to a low-dimensional subspace. In contrast to standard PCA, BiPCA first finds an optimal rescaling of the rows and columns of the data termed *biwhitening* [36]. This rescaling makes the noise homoscedastic and analytically tractable, revealing the rank of the underlying signal. After biwhitening, BiPCA recovers the low-rank signals by removing the transformed noise with optimal denoising techniques [37, 38, 39, 40]. BiPCA is supported by mathematical theory, bridging the gap between previous results in random matrix theory and matrix denoising. Importantly, the pipeline is both verifiable, emitting a goodness-of-fit metric for assessing suitability to a given dataset, and adaptable, alleviating the need to select hyperparameters.

We demonstrate that BiPCA can robustly and effectively estimate the true data dimensionality through simulations and apply it to 123 real datasets from 40 data sources across seven disparate modalities: single cell/nucleus RNA sequencing (scRNA-seq), single cell/nucleus ATAC sequencing (scATAC-seq), spatial transcriptomics, single cell/nucleus methylomics, calcium imaging, single nucleotide polymorphisms (SNPs), and chromatin conformation capture (Hi-C). We demonstrate the importance of choosing the correct rank for downstream single-cell analyses and highlight BiPCA’s ability to reveal the underlying structures in a range of datasets. In addition, we show that BiPCA can greatly improve biological signals compared to other methods, including enhancing marker gene expressions, preserving cell neighborhoods, and mitigating batch effects. BiPCA is available as an open-source Python package at https://github.com/KlugerLab/bipca.

## 2 Results

### 2.1 Overview of Biwhitened Principal Component Analysis

We propose a general, hyperparameter-free, and verifiable pipeline for preprocessing non-negative matrices sampled from a wide range of distributions. Our approach combines our recent method for normalizing random noise matrices [36] with state-of-the-art techniques for recovering low-rank signals from observations contaminated by homoscedastic noise [40]. We couple these techniques to construct a two-step pipeline that recovers a low-rank matrix of latent means from data contaminated by heteroscedastic noise; our pipeline is illustrated in Fig. 1a.

**Figure 1:**
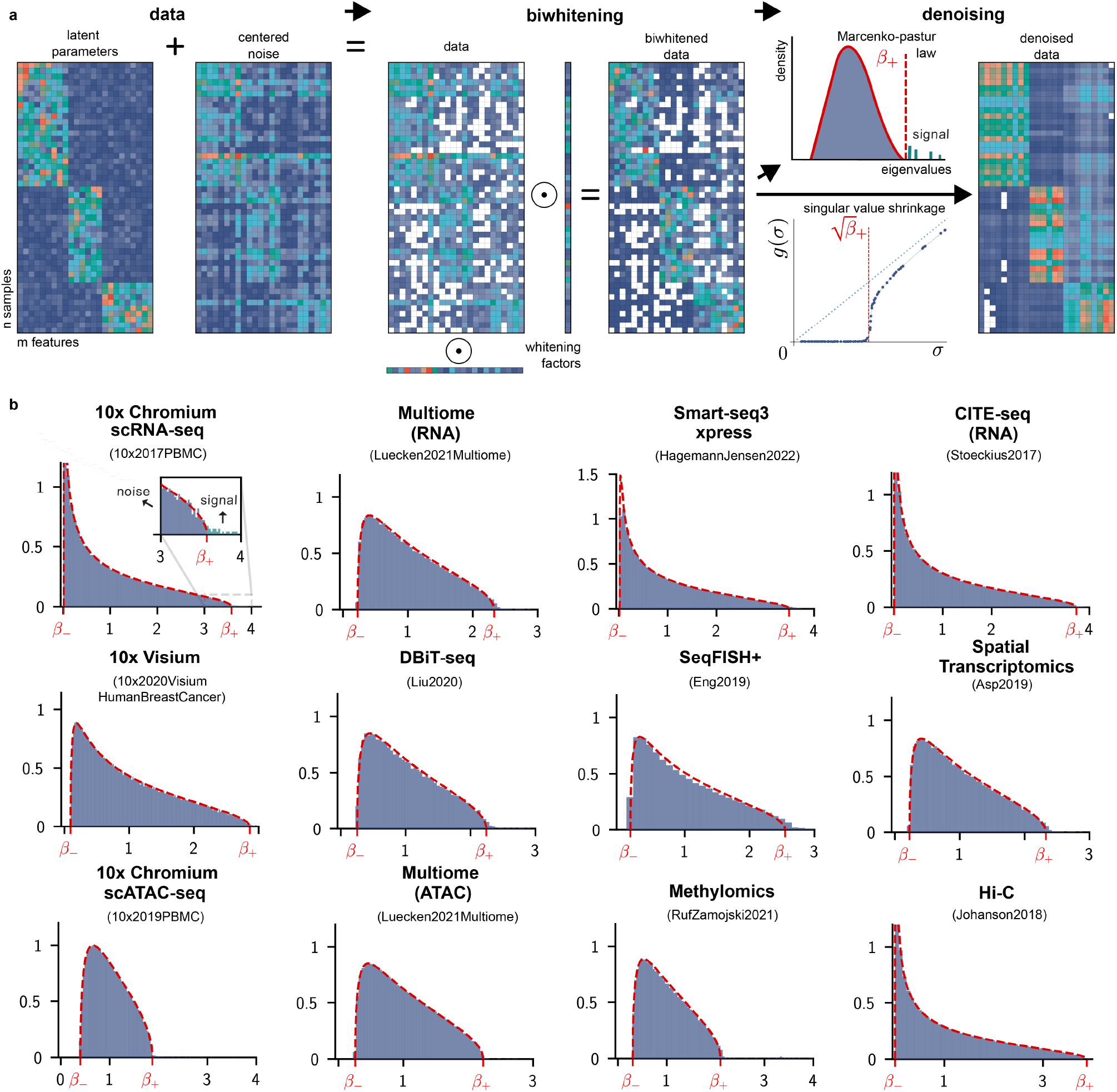
Overview of BiPCA. **a**: BiPCA consists of two steps: *biwhitening* and *denoising*. BiPCA assumes the input count data is the sum of a low-rank matrix of data means and a centered noise matrix (left). Biwhitening (middle) normalizes the rows and columns of the input with *whitening factors*. Biwhitening ensures that the spectral components of the data attributable to noise converge to the Marchenko-Pastur (MP) distribution. Denoising (right) attenuates the effects of noise on the data by removing components with eigenvalues below *β*_+_ (red dashed line in the top panel), or equivalently,removing singular values below 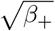 (red dashed line in the bottom panel), and shrinking the singular values above _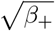_ **b**: Biwhitening is effective for a variety of biological modalities and technologies. Each panel shows that the biwhitened empirical spectral distribution of the data (blue histogram) closely follows the theoretical MP distribution (red dashed curve). The title in each panel indicates the data modality (top) as well as the specific dataset (bottom). _+_ and _*-*_ are the upper and lower edge of each MP distribution. The first panel is zoomed in around _+_ to highlight the separation of signal components from noise.

We model the observed entries of an *m* × *n* data matrix^1^ *Y* as the sum of a low-rank (rank *r* ≪ *m*) mean matrix *X*_*ij*_ = 𝔼 [*Y*_*ij*_] and a centered noise matrix E, that is,

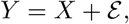

where 𝔼 [ℰ_*ij*_] = 0. This formulation covers a range of count distributions, e.g., *Y*_*ij*_ ∼ Poisson(*X*_*ij*_) where *Y*_*ij*_ is sampled from a Poisson distribution with latent mean *X*_*ij*_. The goal of BiPCA is to *denoise* Y by estimating the latent means matrix *X*.

#### BiPCA extends PCA and singular value shrinkage to the heteroscedastic regime

When a low-rank data matrix *Y* is contaminated by *homoscedastic* noise, i.e. Var[ℰ_*ij*_] = *σ* ^2^ for all entries, the empirical spectral density of the noise covariance *n*^*-*1^ ℰ ℰ^*T*^ can be explained by the Marchenko-Pastur (MP) distribution [35]. In this regime, the spectrum of the data covariance *n*^*-*1^*YY*^*T*^ can be easily split into signal and noise components [36]. As shown in the right side of Fig. 1a, the signal components of the data (i.e., those contributed the signal matrix *X*) will correspond to the leading r eigenvalues of the data covariance *n*^*-*1^*YY* ^*T*^ which are larger than the upper edge of the MP distribution 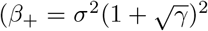, where γ = *m*/*n*), while the density of the noise components (i.e., those contributed by ℰ) will agree with the MP distribution. Equivalently, one can perform this operation directly on the singular value decomposition (SVD) of *Y* by considering the singular values of *n*^*-*1^*Y* larger than 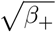.

Using the estimated rank, PCA provides a low-dimensional representation of the data while the closely related truncated SVD provides the closest low-rank approximation to *Y*. Moreover, the quality of this low-rank approximation as an estimator of *X* can be further improved by suitably shrinking the singular values of *Y* [40].

Unfortunately, homoscedasticity is a restrictive requirement for experimental data. For instance, count distributions are common models for biological datasets. Even relatively simple count distributions, such as independent but non-identical Poissons generate data contaminated by heteroscedastic noise [36]. As a result, their spectrum can not be directly modeled by the MP distribution, as shown in Fig. S1 for a range of modalities. The key idea of our approach is to first transform the data such that it satisfies the same spectral properties as homoscedastic noise.

We use *biwhitening*, an algorithm we recently proposed [36]. Biwhitening normalizes the data using row and column *whitening factors û* and 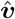,

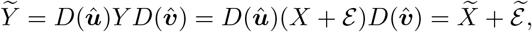

where 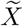 and 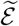 are biwhitened signal and noise matrices, and *D*(***û***) and 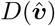 are diagonal matrices of ***û*** and 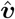. The whitening factors ***û*** and 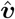 are named so because they are selected to ensure that the average variance is 1 in each row and column of the noise matrix 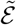 i.e.,

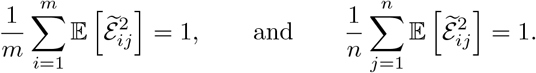

When normalized in this manner, the spectrum of 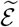 converges to the MP distribution [36]. Because non-zero diagonal scaling preserves the rank of the original unscaled matrix, we can estimate the rank of *X* using the spectrum of 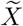, which will be given by the singular values of the biwhitened data matrix 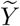 that are not explained by the MP distribution [36].

Once the data has been transformed to this homoscedastic-like regime, the second step of BiPCA is to estimate its biwhitened signal matrix 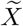 using low-rank approximation. We use singular value shrinkage, which constructs an estimate 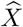 from the spectrum of 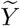,

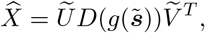

where 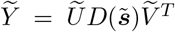, is the singular value decomposition of 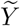 and *g* is an *optimal shrinker* ^2^. The shrinkage function *g* is derived to optimally estimate an underlying signal matrix according to a prescribed loss function. Practically speaking, shrinkage by *g* removes singular values associated with homoscedastic noise, while attenuating signal singular values according to the amount of noise in their corresponding singular vectors. In this work we consider the Frobenius shrinker *g*_*F*_, which minimizes the Frobenius loss 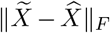 [37], though our package contains implementations of other shrinkers proposed by [40].

After performing biwhitening and singular values shrinkage, BiPCA produces a denoised low-rank matrix that we subsequently demonstrate is well-suited for downstream analysis. However, further postprocessing steps may be required depending on the particular goals of the application. For a more in-depth discussion, we direct the reader to the Methods section.

#### Biwhitening is applicable to a multitude of data distributions

In [36], we showed that the correct whitening factors ***û*** and 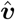 can be learned by solving

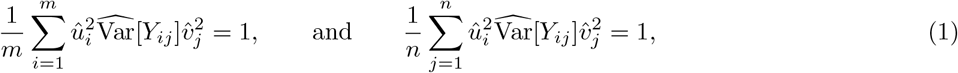

using Sinkhorn matrix scaling [41], where 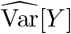 is an unbiased variance estimator, i.e.,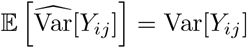.

Conveniently, unbiased variance estimates can be computed directly from the data for a large family of distributions. In particular, if the data *Y*_*ij*_ are sampled from a distribution that admits a quadratic relationship between its variance and mean, namely

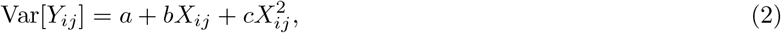

it has a quadratic variance function (QVF) [42]. Distributions of QVFs with *c* ≠ − 1 admit the unbiased variance estimator

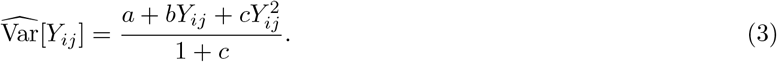

We refer to the distributional parameters *a, b*, and *c* as the intercept, linear, and quadratic coefficients. Distributions with this property include the normal, log-normal, Poisson, generalized Poisson, gamma, binomial with *n* ≥ 2, negative binomial, beta, beta-binomial, and infinitely-many other families of distributions [36].

#### BiPCA is an adaptive algorithm

While it is reasonable to assume that count datasets may be generated by a QVF, the underlying parameters *a, b*, and *c* are typically unknown. To address this, we developed a data-driven strategy to learn quadratic variance coefficients. First, we note that for a given data matrix *Y*, its whitening factors ***û*** and 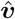 are solely determined by *a, b*, and *c* (Equation (1) and Equation (3)). Thus we can parameterize the biwhitened spectrum of a given matrix by its QVF parameters. Furthermore, since the empirical spectral distribution (ESD) of appropriately biwhitened data converges to the MP distribution, goodness-of-fit for a particular choice of parameters can be measured by using a distributional distance between the normalized data’s ESD and MP law.

Our approach is to minimize the Kolmogorov-Smirnov (KS) distance between the biwhitened ESD and the MP distribution over the domain of *a* = 0, *b* ≥ 0, and *c* ≥ 0. This limits our search space to non-negative count distributions we expect to encounter in real data (besides binomial, which may be specified by the user). In contrast to other works, which estimate mean-variance relationships in data using the empirical mean and variance of each feature [27, 28], our optimization does not assume that the data signal is constant across observations. Our pipeline employs an efficient polynomial approximation-based routine to form estimates 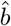 and *ĉ*in a fixed number of iterations, which we describe in our methods.

#### BiPCA is suitable for numerous biological modalities and technologies

To demonstrate the versatility of our adaptive algorithm across contemporary biological modalities, we tested our approach on a compendium of 123 publicly available biological datasets. This repository included seven modalities: scRNA-seq, scATAC-seq, spatial transcriptomics, single cell/nucleus methylomics, calcium imaging, SNPs, and Hi-C. When available, we included datasets sourced from multiple technologies (each with its own capture procedures) for many modalities. Fig. 1b demonstrates the MP fits obtained using KS goodness-of-fit optimization on a subset of these real-world datasets.

### 2.2 Analysis of BiPCA parameter estimates

#### 2.2.1 BiPCA correctly estimates variance parameters and rank in simulations

To assess the performance of our variance parameter estimation algorithm in a controlled setting, we applied our procedure to simulated data of prescribed rank and QVF parameters. Fig. 2a compares ground truth ranks to BiPCA rank estimates 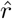 in 5000 × 5000 matrices with Poisson-sampled entries. It is evident that BiPCA robustly recovers the ground truth ranks in all cases. We refer the reader to the supplement section 5.4.1 for more details and discussions of rank estimation accuracy in more challenging regimes.

**Figure 2:**
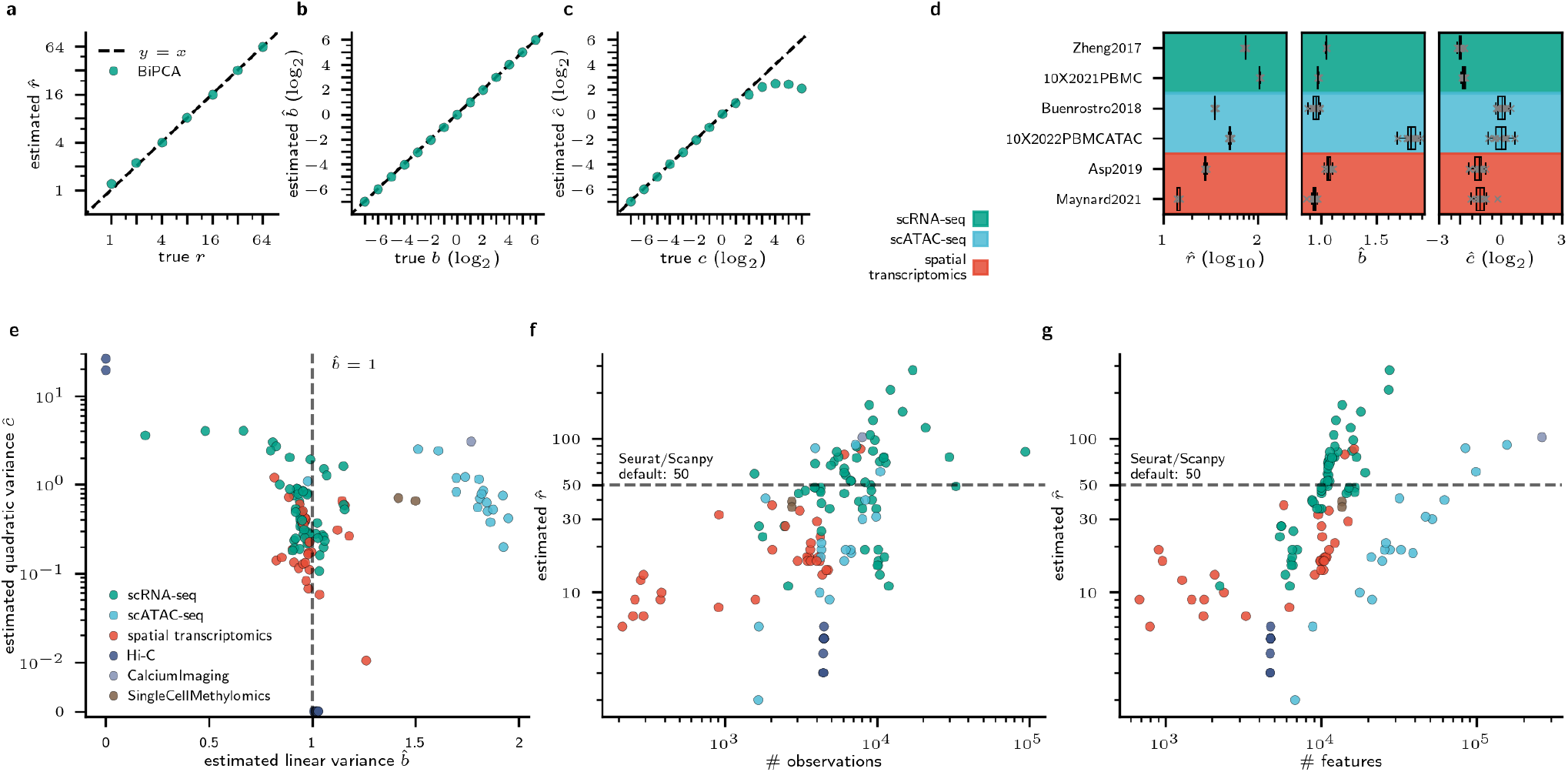
Analysis of BiPCA parameter estimates. **a-c**: BiPCA correctly estimates data ranks r (**a**), linear variance coefficients b (**b**), and quadratic coefficients c (**c**) in simulations. Ground-truth parameters are shown on the x-axis; BiPCA estimates are provided on the y-axis. **d**: Box plot visualization of estimates 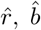 nd *ĉ*(x-axis) across 6 subsampled datasets (y-axis) grouped by modality (scRNA-seq, scATAC-seq, and spatial transcriptomics). Each dataset is resampled 10 times with 75% of the rows from the original data. **e**: Visualization of estimated quadratic variance coefficients 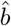 (x-axis) and *ĉ*(y-axis) across surveyed datasets. Each dataset is colored by the corresponding modality. **f-g**: Relationship between the estimated rank 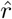 (y-axis) with respect to the number of observations (x-axis in **f**) and the number of features (x-axis in **g**). Datasets are colored by the corresponding modality.

We performed a similar experiment to verify BiPCA’s ability to estimate QVF parameters. Fig. 2b and Fig. 2c compare the ground truth linear and quadratic variance coefficients of simulated 5000 × 5000 rank-1 negative binomial matrices to BiPCA estimates of these parameters. BiPCA provided accurate estimates (y-axis) of the ground-truth QVF coefficients (x-axis) in this simulation, though its ability to estimate *c* became saturated for *c*> 4.

#### 2.2.2 BiPCA reveals the rank and distributional parameters of many biological data modalities

After demonstrating that BiPCA accurately estimates data rank and QVF parameters in controlled simulations, we next aimed to evaluate our algorithm’s parameter estimates across our collection of test datasets.

##### BiPCA reproducibly estimates the rank and distributional parameters in real data

Reproducible parameter estimation is vital for real-world analysis. Here, we studied whether the parameter estimates from BiPCA are consistent with respect to the number of observations in a dataset. We generated distributions of BiPCA parameter estimates obtained from 10 random subsets with 75% of the observations from six real datasets across three modalities (scRNA-seq, scATAC-seq, and spatial transcriptomics). Fig. 2d demonstrates that these distributions exhibit low variance, indicating that BiPCA is robust to variation in data sampling.

##### Biological datasets exhibit a range of generating distributions

After validating that BiPCA’s estimates were consistent across subsets of the same data, we further examined inter-experiment variances of the parameter estimates. We plot the *b* and *c* estimates across a range of datasets colored by modalities in Fig. 2e. Furthermore, we concentrate on scRNA-seq, spatial transcriptomics, and scATAC-seq, the three most abundant modalities in the compendium, to explore the variations across different protocols within each modality, as shown in Fig. S2a-c.

We observed that the estimated linear variance coefficient of most scRNA-seq and spatial transcriptomic datasets was approximately 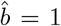 (Fig. 2e, Fig. S2a-b), whereas the estimated quadratic variance coefficient *ĉ*exhibited a range of positive values, though most were less than 1. Together, this suggests that the variance estimates from BiPCA on these modalities fall under negative binomial distributions (*b* = 1 and *c* > 0). On the other hand, most of the 10x datasets we analyzed had similar parameter estimates (Fig. S2a); however, we did find that Smart-seq datasets exhibited a broad range of linear variance and higher quadratic variance in comparison to 10x.

Surprisingly, most scATAC-seq datasets we studied exhibited larger linear variance coefficients than a standard negative binomial (Fig. 2e). The mean 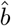 of these datasets was 1.797. Closer inspection revealed that the datasets with large linear variances were all generated using 10x scATAC-seq protocols (10x Chromium V1 and 10x Multiome). These protocols use read counts from paired-end reads as count values. It has been shown that paired-end read counting double counts ATAC data, leading to variance almost twice as large as the mean [13]. One proposed alternative is to compute fragment counts by rounding all uneven counts to the next highest even number and halving the resulting read counts [13]. When reanalyzed using this fragment counting procedure, the mean estimated linear variance coefficient of 10x ATAC datasets was 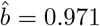 Despite the differences in terms of count distributions, we found no substantial differences in terms of the estimated rank between the two counting strategies (Fig. S2d). We also analyzed data collected using the protocol of the original scATAC-seq work [43], which used fragment counts as count values in their pipeline. Interestingly, the linear variance coefficient estimate for this data was 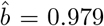, which closely matched the parameters we estimated after applying fragment counting to 10x data. In summary, we find that the paired-end read counting strategy used by 10x protocols may induce extra linear variance, but when fragment counting is used, BiPCA estimates that scATAC-seq data is closer to a negative binomial distribution. Next, we surveyed the estimated data rank across our repository. We evaluated the estimated rank with respect to the throughput (the number of observations) and the number of features. As shown in Fig. 2f and g, the rank varies dramatically between datasets. As expected, rank is positively correlated with the number of observations and features both within each modality and across modalities. This indicates that signal complexity grows when improving throughput and coverage in general. In addition, we observe rank patterns that are specific to different modalities. For instance, spatial transcriptomics exhibited lower estimated ranks across datasets, likely due to intrinsic cell or gene throughput limitations across spatial technologies. On the other hand, some datasets have higher rank estimates, indicating more complex signal structures. Notably, BiPCA estimated that 52.8% of the scRNA-seq datasets we considered have a higher rank than 50, which is the default number of principal components to use when performing PCA using Seurat [44] and Scanpy [45] on scRNA-seq data. In **Section** 2.3.1 we explore the impact of improper rank selection for data analysis and highlight the importance of using the correct rank.

### 2.3 BiPCA enhances downstream single cell analyses

In the ensuing results, we demonstrate BiPCA’s improvement of single cell analyses in comparison to other approaches. For scRNA-seq datasets, we compared BiPCA with five methods: log-normalization (log1p), log-normalization with gene-wise standardization (log1p+z), Pearson residuals (Pearson) [28], Sanity [29], and ALRA [46]. In scATAC-seq datasets, we compared BiPCA with three normalization methods: log-normalization (log1p), log-normalization peak-wise standardization (log1p+z), and Term Frequency -Inverse Document Frequency (TF-IDF) [47].

#### 2.3.1 Proper rank estimation is critical for exploratory analysis of single cell data

Dimensionality reduction using 50 principal components (PCs) has become the de facto standard for single cell and spatial transcriptomic data analysis; it is the default rank used by Scanpy [45] and Seurat [44]. Alternatively, one can use scree plots to select rank based on cumulative variance, but this approach does not differentiate between variance from the signal and variance from the noise. BiPCA, in contrast, reveals the true rank of the data by providing a spectrum that can be easily partitioned into signal and noise using the MP law. In this section, we show that proper rank selection is crucial, as improper choices can bias and degrade exploratory data analysis. We focus on two situations: first, when the rank is *overestimated* and second, when the rank is *underestimated*.

##### Rank overestimation masks structure in data with low signal-to-noise ratio (SNR)

In low SNR settings, the magnitude of the leading r eigenvalues of the data covariance are small; when one selects too high of a rank, they consequently retain noise components whose total magnitude is close to the magnitude of the retained signal. This noise can obscure one’s ability to recognize structure in the data, such as the clustered Poisson simulation shown in Fig. 3a-b. This simulation is composed of five clusters (in analogy to single cell data composed of multiple cell types), with each cluster drawn from a single linear independent signal component (see Methods for details). If one were to only analyze the t-SNE visualization of the rank 50 approximation of this data (Fig. 3a) in the absence of our ground truth labeling, they might conclude that there is no cluster structure. On the other hand, t-SNE visualization of its rank five approximation (Fig. 3b) reveals the data’s underlying structure: five separate clusters.

**Figure 3:**
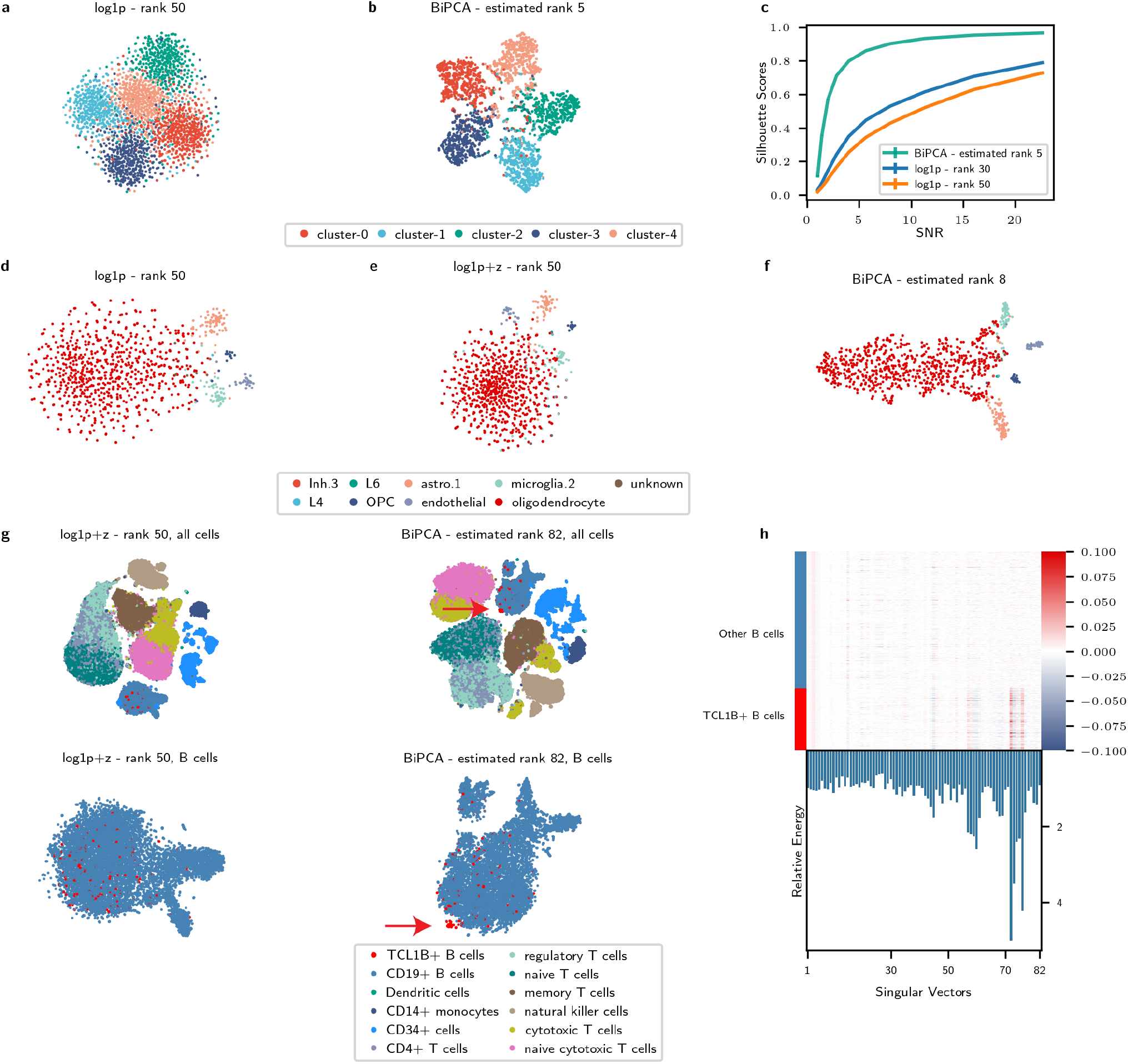
Proper rank estimation is critical for downstream analyses. **a-b**: t-SNE visualization of a simulated rank five data using rank 50 approximation (**a**) and rank five approximation (**b**). Cells are colored by the ground truth cluster labels. **c**: Comparison of silhouette scores between rank 30 approximation / rank 50 approximation / BiPCA rank approximation across different SNR regimes on the simulated data. **d-f** : t-SNE visualization of the cosMx human frontal cortex data (one FOV) using rank 50 approximation from log1p (**d**) and log1p+z (**e**), and rank eight approximation from BiPCA (**f**). **g**: t-SNE visualizations of Zheng2017 PBMC data using different normalization methods and different number of signal components for low rank approximation. Top left: log1p+z normalization with rank 50 approximation. Top right: log1p+z normalization with rank 82 approximation. Bottom left: zoom-in visualization of the B cells from the top left embeddings of log1p+z. Bottom right: zoom-in visualization of the B cells from the top right embeddings of BiPCA. Cells are colored by the cell type labels, and red arrows highlight the TCL1B+ B cell population. **h**: Top: heatmap visualization of the signal components (singular vectors) across B cells for the Zheng2017 PBMC data. The columns are singular vectors ordered by singular values, and the rows are B cells group by B cell labels. Bottom: Relative energy of the TCL1B+ B cells within all B cells for each singular vector. The singular vectors with the largest relative energy are indexed at: 72, 73, and 76.

We used this simulation to analyze the minimum SNR required to recover the underlying cluster geometry of the data using rank 30 approximation, rank 50 approximation, and BiPCA. As shown in Fig. 3c, BiPCA achieves superior silhouette scores compared to other approximations across SNRs from one to ≈ 20. Notably, the gap in performance between BiPCA and fixed approximation methods is greatest for low SNRs, and as the SNR exceeds 10, BiPCA’s silhouette plateaus close to optimality (one) when SNR is large. In addition, at each SNR we studied, BiPCA successfully recovered the correct rank (five). This experiment demonstrates that over-estimation of the rank in low SNR settings can obscure data geometry. BiPCA, which is effective even in low SNR regimes, avoids this situation by adapting to the data.

The perils of rank overestimation are not exclusive to toy examples. In Fig. 3d-f, we focus on one field of view (FOV) from a human frontal cortex CosMx Spatial Molecular Imager (SMI) dataset [48]. BiPCA estimated 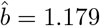 and *ĉ*= 0.264 for this data, and the recovered rank was eight. Fig. 3d-f compares the t-SNE visualizations of log1p+z with 50 PCs, log1p+z with 30 PCs, and BiPCA with eight PCs. As shown in Fig. 3d-f, different cell types are mixed together when using an excessive number of PCs (e.g., 50 and 30, with silhouette score of 0.03 and 0.129, respectively). However, they form clear cluster structures when the correct number of PCs is used (rank eight, with silhouette score of 0.336).

##### Rank underestimation ignores fine structures in scRNA-seq data

When data rank is underestimated, weaker signal components are excluded from downstream analysis. These components represent orthogonal sources of variation that can capture fine structures in the data. These structures may arise from biologically relevant phenomena such as rare cell types or minor perturbations in cell state. To illustrate this, we analyzed the entire *Zheng2017PBMC* dataset [49], which is composed of 10 cell types of purified peripheral blood mononuclear cells (PBMCs). The estimated rank and QVF parameters of this data were 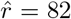, 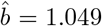and *ĉ*= 0.272. Analyzing this data using BiPCA revealed a subset of CD19+ B cells that uniquely express TCL1B (Fig. S3a). TCL1B expression is enriched in naive B cells [50]. Critically, TCL1B+ B cells were indistinguishable when one considered rank 50 and/or log1p preprocessing of the data (Fig. 3g). However, they formed a distinct and separable cluster in the BiPCA denoised data after including all 82 signal components. To confirm this separation is captured by trailing signal components, we inspected the values of the normalized data’s singular vectors over the B cells (Fig. 3h). Several trailing singular vectors (e.g., No. 72 and 76), clearly distinguished TCL1B+ B cells from other B cells. This was further confirmed by the large energy of these singular vectors within the TCL1B+ B cells in comparison to all B cells (Fig. 3h, bottom).

To further demonstrate that BiPCA can identify novel cell populations in complex systems, we applied BiPCA to the RNA modality of one 10x Multiome human prefrontal cortex sample from the SCORCH consortium [51]. BiPCA estimated the rank of this data as 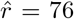. We identified a subset of Oligodendrocyte precursor cells (OPCs) with unique SEMA3E expression (labeled as SEMA3E+ OPCs). These SEMA3E+ OPCs have signals strongly enriched in component No. 55 and 56 as shown in Fig. S4a, which would be ignored by standard pipelines that use only 50 components. Besides SEMA3E, this group of SEMA3E+ OPCs also shows other unique marker expression profiles (Fig. S4b). We repeated this experiment for two other samples in this cohort and found that weak signal components were critical for recovering this subpopulation (Fig. S4c-f) across samples.

#### 2.3.2 BiPCA enhances biological signals in single cell data

Our next objective was to evaluate the quality of biological signals extracted by BiPCA.

##### BiPCA enhances marker gene coherence in scRNA-seq

Marker genes are often employed in scRNA-seq for supervised cell annotation because they are typically well-characterized and reproducible indicators of phenotype. To evaluate whether BiPCA is effective for extracting biological signals, we assessed the coherence of normalized marker gene expression with the cell types they are known to identify. This experiment required two external sources of ground truth: a set of ground truth cell type labels (ideally identified without scRNA-seq), and a set of canonical mappings between genes and the cell types they mark. For the former, we compiled an experimental dataset, *Zheng2017-markers*, composed of the flow-sorted populations of CD4+ T cells, CD8+ T cells, CD19+ B cells, and CD56+ natural killer (NK) cells from the *Zheng2017* dataset [49]. For the latter, we extracted a set of 101 marker gene annotations using the Human Protein Atlas (HPA) [50]. We used *Zheng2017-markers* and these curated marker genes to quantitatively evaluate how well each marker gene separated its annotated cell type following transformation with log1p, log1p+z, Pearson, Sanity, ALRA, and BiPCA. BiPCA estimated that the QVF parameters of this dataset were 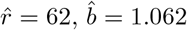, and *ĉ*= 0.081.

We first focus on four prominent marker genes, CD40, NCAM1, CD4, and CD8A. CD40 is a well-characterized B cell marker, while the protein products of NCAM1 (synonymous with CD56), CD4, and CD8A were used during the flow sorting that generated our reference dataset; thus, we expected that these RNA markers should be effective one-dimensional linear classifiers for their targeted cell type. Instead, we found that normalization with log1p, log1p+z, Pearson, and Sanity was ineffective for separating targeted cell types from background cell types using marker genes (Fig. 4a). For these methods, the bulk of normalized gene expression for both targeted and untargeted cell types was concentrated near zero, and the resulting area under the receiver operating characteristic curve (AUROC) was small (≤ 0.71). On the other hand, marker gene classifiers that used low rank approximation (ALRA and BiPCA) achieved perfect classification of B cells and NK cells. Notably, while transformation with ALRA yielded good CD8+ and CD4+ T cell classifiers (AUROC 0.92 and 0.97, respectively), BiPCA achieved perfect or near-perfect classification of all cell types we surveyed.

**Figure 4:**
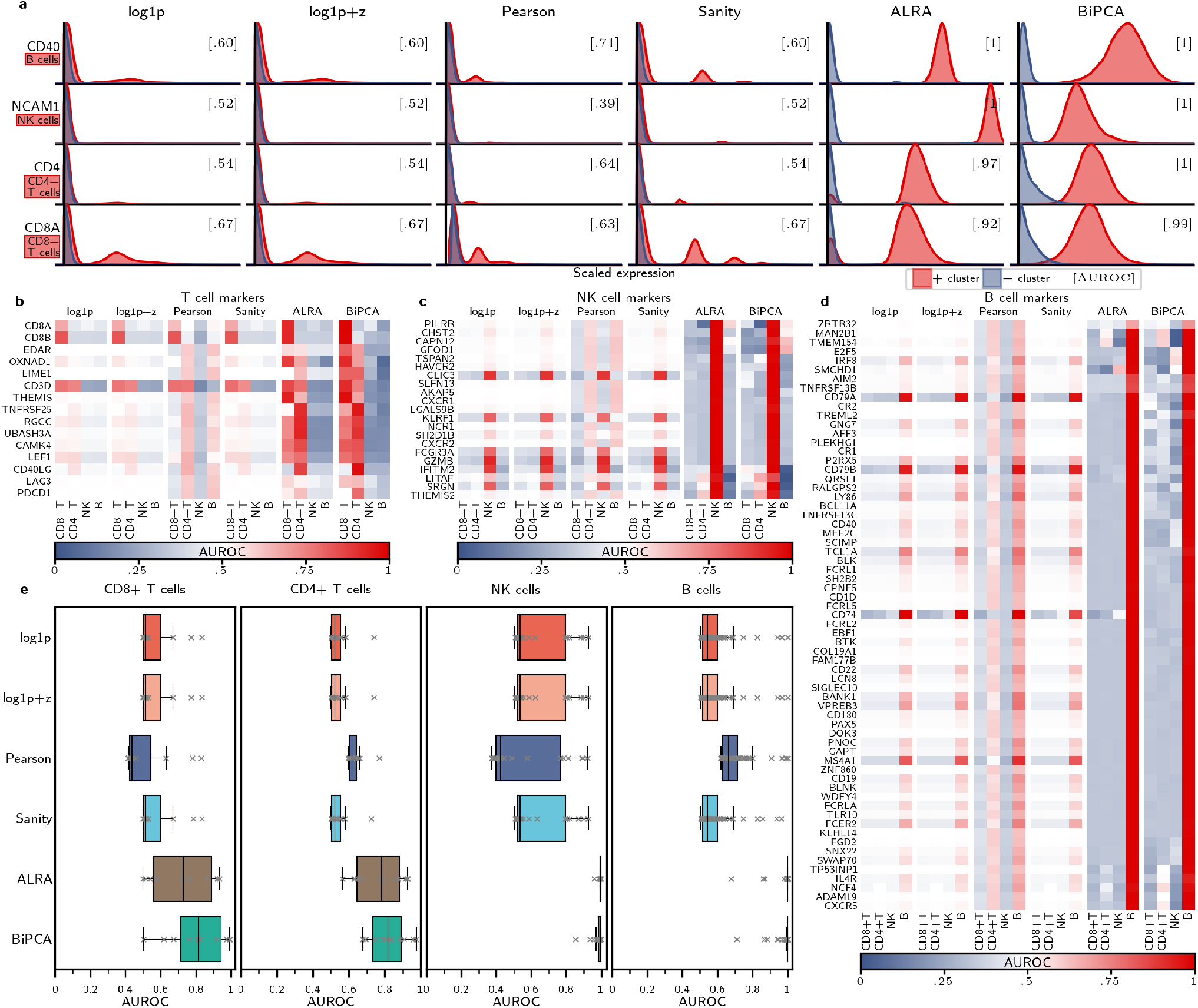
Comparison of marker gene expressions across normalization schemes using *Zheng2017* dataset. **a**: comparing the distribution of 4 marker genes (labeled on the rows, together with the associated cell type) across different normalization methods (columns). Each red distribution represents the scaled expression level of the cluster associated with the marker, and each blue distribution represents the scaled expression level from all the other clusters. AUROC values are computed by classifying the two distributions (shown in brackets). **b-d**: Heatmap visualizations of the classification AUROC values using marker genes associated with T cells (**b**), NK cells (**c**), and B cells (**d**) from the Human Protein Atlas. The marker genes are shown on the rows and the cell types are shown across columns split by normalization methods. AUROC values are computed by classifying each cell type against others for each marker gene. **e**: Box plot visualization of the AUROC values for different markers associated with each cell type (columns) across different normalization methods (rows).

We hypothesized that the properties of low-rank approximation would induce strong correlations between markers for specific cell types, thus enhancing the signal of less prominent marker genes in BiPCA and ALRA in comparison to the other approaches we compared. To study this, we repeated our previous one-dimensional linear classification experiment using the list of putative markers we extracted from the HPA. Fig. 4b-d report the classification AUROCs for 15 T cell markers, 21 NK cell markers, and 65 B cell markers respectively. In general, ALRA and BiPCA outperformed other methods at this task, and the distributions of AUROCs (Fig. 4e) reflected this. However, we note that BiPCA performed best at this task on CD4+ and CD8+ T cells with median AUROCs exceeding ALRA, demonstrating stronger ability to enhance marker coherence.

Induction of false correlations between cell types and gene expression is one concern for low rank approximation of scRNA-seq [29]. In our classification experiment, false correlations would manifest as AUROCs greater than 0.5 for off-target cell types. We observed no such off-targets in T cell marker genes normalized using BiPCA. BiPCA returned eight candidate off-targeting markers from NK cell-marking genes: PILRB, CHST2, CAPN12, and GFOD1 exhibited small positive AUROCs for B cell classification; IFITM2 was a weak marker for CD8+ T cells; and LITAF, SRGN, and THEMIS22 were weak markers for CD4+ T cells. Similarly, we observed six candidate off-targeting B cell-markers: MAN2B1, TMEM154, NCF4, and CXCR5 were weak classifiers of CD4+ T cells; while E2F5 and IRF8 weakly encoded for NK cells. Although each marker is annotated by the HPA as enriched in either NK or B cells, closer inspection of the HPA confirmed that elevated expression of these genes relative to other cell types has been observed in the cell types for which we observed AUROCs greater than 0.5. Thus, we contend that supposed false correlations between marker gene expression and flow-sorted cell type labels after applying BiPCA are biologically plausible.

##### BiPCA improves biologically meaningful embedding of cell neighborhoods

A foundational assumption of single-cell biological analyses is that cell neighborhoods reflect underlying biology [52]. When this assumption holds, cell clusters and visually separable cell groups in low-dimensional neighbor embeddings may be interpreted as distinct phenotypes. We sought to quantify whether BiPCA preserved or improved biological information in its PCA space relative to other normalizations by evaluating neighborhood embedding on three dataset collections: (1) CITE-seq (RNA): the RNA modality of 14 CITE-seq datasets including *Stoekius2017* [53], *Stuart2019* [54], and 12 batches from *Luecken2021CITE* [55], (2) Multiome (RNA): the RNA modality of 13 10x Multiome datasets (13 batches from *Luecken2021Multiome* [55]), and (3) Multiome (ATAC): the ATAC modality of the same 13 10x Multiome datasets. Many metrics for measuring embedding quality with respect to underlying biology have been proposed [56]. In Fig. 5 we consider the silhouette coefficient, which measures how compact and separable labeled cell types are relative to each other in a given embedding: larger values indicate that cell types are more separable and tightly clustered in an embedding. As shown across datasets in each modality (CITE-seq (RNA) in Fig. 5a, Multiome (RNA) in Fig. 5b, and Multiome (ATAC) in Fig. 5c), BiPCA consistently improved the silhouette score of ground-truth cell labels, illustrating its superior performance to recover tight cell neighborhoods. In addition, we also computed k-nearest neighbor (kNN) accuracy that measures the how well each method preserves the neighborhood such that the nearest neighbors are from the same labeled cell type (Fig. S5). BiPCA performs at par with other normalization methods in the CITE-seq (RNA) datasets and Multiome (RNA) datasets and consistently outperforms other normalization methods in the Multiome (ATAC) datasets. This illustrates that BiPCA preserves cell neighborhoods with respect to labeled biology.

**Figure 5:**
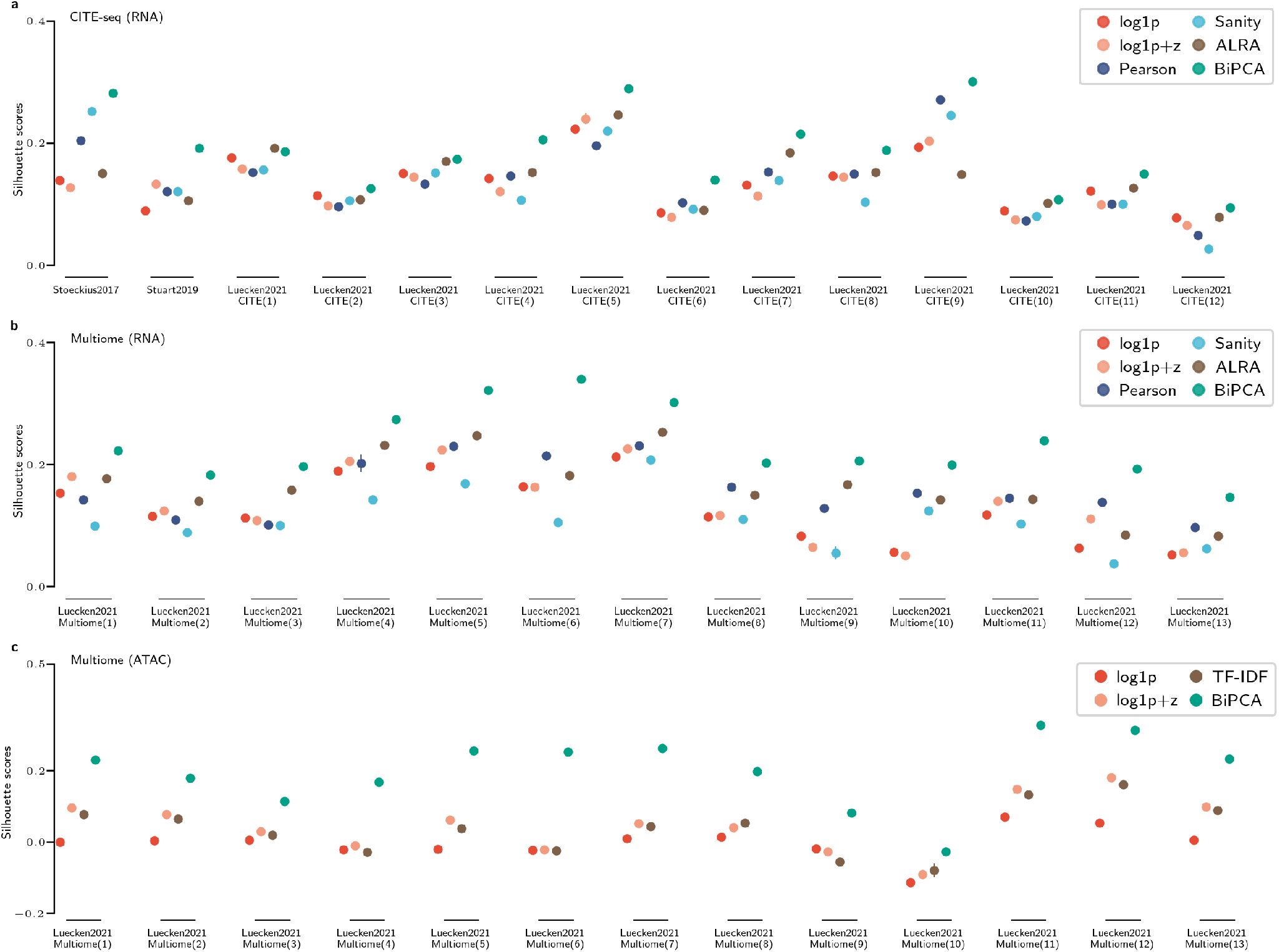
Comparison of silhouette scores with respect to the cell types across 3 sets of datasets (**a**) CITE-seq (RNA): the RNA modality of 14 CITE-seq datasets (**b**) Multiome (RNA): the RNA modality of 13 Multiome datasets (**c**) Multiome (ATAC): the ATAC modality of the same 13 Multiome datasets as in **b**. Each experiment is repeated 10 times with 80% of the data. Within each dataset, different normalization methods are applied (colored dots) and silhouette scores are computed on the PCA space in each method. Mean silhouette scores and the standard deviations are plotted.

#### 2.3.3. BiPCA removes sources of confounding noise in single-cell experiments

Due to the stochastic nature of the read sampling process, single-cell data is often plagued with sampling noise that confounds the underlying biological variance. For example, [27] demonstrated that differences in cell sequencing depths can dramatically affect downstream analyses. We hypothesized that these effects could be mitigated using biwhitening. To evaluate this, we applied BiPCA to a 10x Chromium single-nucleus RNA sequencing (snRNA-seq) dataset from a human insular cortex sample [51]. In this dataset, four technical replicates were sequenced from the same sample. Dramatic library depth differences between replicates were evident in this data (Fig. S6).

Fig. 6a compares t-SNE visualizations produced after normalization of this dataset. We observed clear batch effects in which regions of different clusters are dominated by one replicate. In particular, cells from replicate one tended to embed separately from cells of the other replicates under all normalizations other than BiPCA. The difference in replicate density was highly evident when we focused on astrocytes (Fig. 6b).

**Figure 6:**
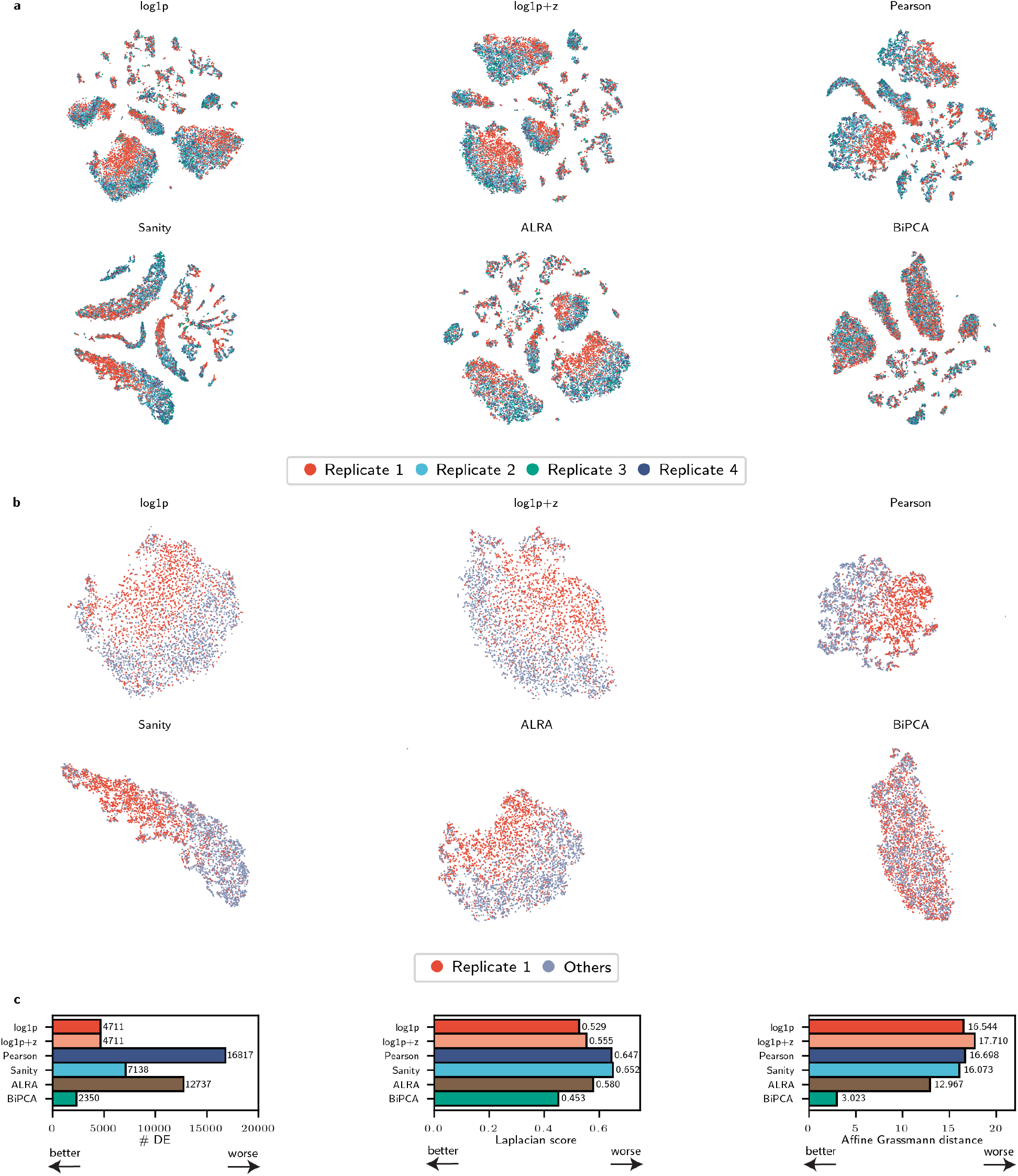
BiPCA mitigates batch effect across technical replicates of a 10x Chromium single-nucleus RNA sequencing (snRNA-seq) dataset from a human post-mortem insular cortex sample. **a**: t-SNE visualization of different methods on the whole dataset. Cells are colored by the replicate ids. **b**: Zoom-in t-SNE visualization of the astro-cyte sub-population. Cells are categorized and colored as replicate 1 and other replicates. **c**: Comparison of the performance of batch effect correction between different methods measured by 3 different metrics. Left: number of differentially expressed genes (# DE), middle: Laplacian score, right: affine Grassmann distance. x-axis shows the values of each metric and y-axis shows different methods. In all metrics, smaller values represent better performance, while larger values represent worse performance.

To further characterize the drivers of differential technical replicate embedding, we compared astrocytes from replicate one to all other replicates using three metrics (Fig. 6c). First, we examined the number of differentially expressed genes (# DE) between replicate one and the other replicates (Fig. 6c, left). We expected # DE to be small for technical replicates; however, we found that no normalization returned zero DE genes. On the other hand, BiPCA returned the fewest differentially expressed genes (# DE = 2350), a two-fold improvement over the next best methods (log1p and log1p+z, # DE = 4711). Next, we computed the Laplacian score [57] of the replicate one indicator vector on a cell-cell graph (Fig. 6c, middle). This metric measures the smoothness of the replicate labels; we expected that technical replicates should be well-mixed, and thus their label distribution would be high frequency with respect to the underlying cell-cell graph structure, leading to low Laplacian scores. BiPCA performed the best at this metric. Finally, we measured the alignment between the post-normalization subspaces spanned by replicate one and the other replicates using the affine Grassmann distance [58, 59]. We expected that technical replicates should in general span well-aligned subspaces if there are no significant replicate effects, and thus the corresponding affine Grassmann distance would be small. BiPCA achieved the lowest score in this metric, indicating that the post-normalized replicate subspaces captured similar structure. In conclusion, numerical quantification indicates that BiPCA was effective at mitigating replicate effects introduced by sampling bias.

In addition to snRNA-seq, we also investigated BiPCA’s ability to attenuate the impact of library depth differences in scATAC-seq analyses. While normalizations such as log1p and TF-IDF are widely adopted for scATAC-seq preprocessing [60, 47], the effect of library depths still manifests strongly in the leading components of normalized data [60]. We inspected the principal components of the ATAC modality of *Luecken2021Multiome* after applying BiPCA for evidence of confounding library depth signals (Fig. S7). Principal components after BiPCA were significantly less correlated with library depth in comparison to log1p, log1p+z, and TF-IDF. Together with the results shown in Fig. 6, this demonstrates that BiPCA can improve downstream analyses by reducing the impact of library depth biases on biological signals.

#### 2.3.4 Variance stabilization and stable low-rank approximation in scRNA-seq

Finally, we sought to evaluate the effects of BiPCA on the mean-variance relationship of features in single cell data. During these experiments, we discovered that many methods exhibit dramatic fluctuations in gene variance which could affect reproducibility.

##### Biwhitening stabilizes gene variance with respect to gene mean

Fig. S8a plots the relationship between gene mean and gene variance in the raw UMI counts of a 33092 cell × 11055 gene reference dataset, *10×2016PBMC* (also referred to as *pbmc33k* by [27]). The evident heteroscedasticity in this dataset is prototypical of scRNA-seq experiments. The aim of variance stabilization (through, e.g., the log1p transform, Pearson residuals, or other methods) is to control for heteroscedasticity in the data; intuitively, one seeks a transformation in which the variance of a gene is proportional to its biological variability, rather than its abundance.

We applied log1p transform, Sanity, Pearson residuals, and BiPCA to *10×2016PBMC*. BiPCA’s estimated parameters were 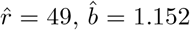, and *ĉ*= 0.592. Fig. S8b-e show the transformed gene-wise variances as a function of the gene’s original mean using the log1p transform, Sanity, Pearson residuals, and biwhitening (before denoising) to the original data. In each of these transformations, no low rank approximation was performed. We note that of the four transformations, biwhitening and Pearson residuals perform the best at removing the relationship between the mean and transformed variance. Between these two transformations, Pearson residuals exhibits a much broader range of variance (approximately two orders of magnitude) than biwhitening. The authors of Pearson residuals claim that this residual variance reflects underlying biology [27]. Large ranges in transformed variances can be a useful property for feature selection [28]. However, we note that it has been shown that highly variable gene selection is not essential for achieving state of the art performance in cell type classification [28, 46]. On the other hand, the transformed gene variances after biwhitening are supported on a much narrower range (approximately one order of magnitude). This finding is evidence of the heteroscedastic normalization inherent to biwhitening.

##### BiPCA, standardized log1p, and Pearson residuals preserve variances in lowly-expressed genes

Next we compared transformed gene variances in *10×2016PBMC* using pipelines that perform low rank approximation on the data. These pipelines were rank 50 approximation of the raw data, log1p, log1p+z, Pearson, and Sanity (Fig. S9a-f), along with rank 73 approximation of ALRA (Fig. S9e) and rank 49 approximation of BiPCA (after denoising) (Fig. S9g). Rank 50 approximations were performed as 50 is the default rank in Scanpy [45] and Seurat [44], whereas the rank estimated by ALRA and BiPCA are 73 and 49, separately. Of the considered transformations, only rank 50 approximation of log1p+z, Pearson residuals, and BiPCA exhibited large transformed variances in genes with low numbers of observed counts, demonstrating their ability to recover proportionally large gene variances independent of their original mean.

##### BiPCA recovers stable representations of gene variance

A key assumption for applying PCA to scRNA-seq experiments is that one can robustly estimate the principal axes of variation of the latent biological system by sampling only a subset of the cells during sequencing. Yet, theoretical works suggest that performing PCA on data corrupted by heteroscedastic noise can lead to poor estimation of both singular values and axes of variation [61]. Given the evident heteroscedasticity of scRNA-seq data, we sought to quantify the ability of different preprocessing methods to stably represent scRNA-seq experiments.

Since it is impossible to know the ground truth axes of variation in real data, we cannot quantify whether a pipeline robustly estimates ground-truth principal components. On the other hand, we can quantify how stably a pipeline applied to a subset of the data recovers similar results to the full data. This, in effect, allows us to simulate the regime we desire, in which ground truth biology is known (in this case, the gene variances) but only a small subset of cells are observed (as in a sequencing experiment). To that end, we applied each normalization pipeline to subsets of 5000 cells selected uniformly at random from *10×2016PBMC*. We compared the post-PCA gene variances from each transformed subset of the data to the post-PCA gene variances computed from the transformed complete data (Fig. S10). We expected that normalizations with stable signal reproducibility would generate similar gene variance distributions between the full data and its subsets; this would be indicative that the normalization retains the relative variance between genes even in small experiments. We measured this using the spearman correlation coefficient *r*_*s*_. We interpret low values in this metric to imply that uniform random cell subsampling is sufficient to distort the relationship between biological signals under a normalization. Only BiPCA (*r*_*s*_ = 0.92) and log1p (*r*_*s*_ = 0.97) achieved spearman correlations larger than 0.9. We expected such performance from log1p, as it features no gene-wise transformations and thus expression values do not change under sub-sampling. However, we have shown throughout this work that log1p is consistently outperformed in other aspects by normalizations that feature gene-wise transformations. This experiment shows that those methods (besides BiPCA) are sensitive to sampling, and may produce distorted variance relationships that are difficult to reproduce between samples of the same underlying system. From this we conclude that BiPCA offers superior reproducibility in comparison to other normalizations.

## 3 Discussion

In this study, we presented BiPCA, a principled pre-processing pipeline combining normalization and dimensionality reduction tailored to high-throughput count data. BiPCA first scales the rows and columns of the data to make the noise approximately homoscedastic (*biwhitening* step), thereby revealing the underlying rank of the data (based on random matrix theory). Then, BiPCA performs optimal shrinkage of singular values to recover the biological signal (*denoising* step). BiPCA is an adaptive algorithm that can be applied to a variety of data-generating count distributions, such as Poisson or negative binomial, by virtue of the data-driven quadratic variance estimation procedure. Through simulations, we demonstrated the high fidelity and robustness of BiPCA for fitting diverse data-generating processes and for rank recovery. Using a collection of 123 datasets from seven different modalities, we showed that BiPCA is highly versatile and BiPCA representations preserve and enhance biological signals while removing sampling noise and are useful for downstream analysis.

The optimality of BiPCA relies on several assumptions, most importantly the low-rank structure of the underlying signal and the independent count noise observations. When the low-rank assumption is violated, it becomes difficult to separate the signal spectrum from the noise spectrum (as given by the Marchenko-Pastur distribution) for accurate signal recovery. However, by analyzing 123 real-world high-throughput count datasets, we showed that in practice this assumption is often met; the highest estimated rank that we observed was 280. Though 280 is still a low rank for the dataset with over 27 000 features, it was much higher than the default number of retained principal components (typically 50) in standard analytical pipelines [44, 45].

Another assumption behind BiPCA is that the variance of the true data-generating count distribution is a quadratic function of the mean. We demonstrated that this framework is sufficiently flexible to model many different data modalities; however, we also encountered datasets that were not suited for BiPCA. In such cases, our built-in goodness-of-fit metrics highlight that the dataset does not meet our assumptions and BiPCA is not applicable. For example, in single-cell methylomics, the quantity of interest is the ratio of two counts, methylated sites and total sequenced sites. While biwhitening could appropriately normalize both the methylation count matrix and the total count matrix, applying BiPCA to the entry-wise ratio of these matrices did not yield acceptable fits to the MP distribution. In [62], we show that it is possible to biwhiten such matrices without estimating their variance by computing their entire singular value decomposition; this presents a future opportunity to extend our pipeline.

Quadratic mean-variance relationships encompass a broad range of count distributions such as Poisson, negative binomal, and gamma distributions, offering greater flexibility compared to existing normalization methods, which often rely on restrictive assumptions about the underlying distributions [22, 23, 24, 25, 26, 27, 28, 29]. For scRNA-seq data sets, BiPCA yields variance estimates close to the negative binomial distribution (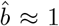, Figure 2e) with substantial overdispersion compared to Poisson (0.1 ≤ *ĉ*≤ 1). Recent work argued that the measurement noise in scRNA-seq data is close to Poisson [28, 63, 29] with *c* ≈ 0. The BiPCA estimates may include not only measurement noise but also some non-low-rank biological variability (biological noise). Importantly, BiPCA does not require any assumptions on the overdispersion strength and instead fits it to the data.

Several recent works have applied random matrix theory to single-cell RNA-sequencing data analysis [30, 31, 33, 32, 34]. These methods all attempt to fit the MP law via different heuristic normalizations. However, most datasets do not conform to the MP distribution after simple library normalization [31] or log normalization with standardization [34]. After log-transformation, [30] fitted the aspect ratio γ of the MP distribution instead of using the actual aspect ratio of the count matrix γ= m/n. Subsequent works pointed out problems with both the log-transformation and aspect ratio fitting [32, 33], and suggested an additional single step of row and column standardizations. [32] demonstrated that this could be used for measuring clusterability, and scLENS [33] proposed this technique for normalization. However, the MP fits produced by these methods are often still unsatisfactory (see, e.g., Figure S2D of [32]). Recently, [34] approached the problem by replacing the MP distribution with a random matrix theory model specifically developed for scRNA-seq data, but their approach only works in the presence of baseline and perturbation measurements.

Our method is different in several key ways. First, BiPCA is fully data-driven. It fits the spectrum of the data to the MP law with the actual aspect ratio after biwhitening, which provides guaranteed convergence and fit [36]. Coupled with the optimal denoising step, our model is more rigorous and robust in removing noise and recovering the signal. Second, our model is highly adaptable to diverse data modalities and complex variance structures, going beyond the scRNA-seq modality. In addition, we do not impose any assumptions on the data due to clustering or differential expression analysis.

The pipeline proposed in this work builds on the biwhitening framework we introduced in [36]. While [36] primarily focused on the analytical aspects of biwhitening, laying its theoretical foundation and highlighting its advantages for rank estimation, this study substantially extends the framework in several important directions. First, we develop an efficient and scalable method for estimating data variance parameters, enabling the analysis of large data matrices. Second, we incorporate singular value shrinkage to denoise the data and recover underlying biological signals, which is critical for downstream analytical tasks beyond rank estimation. Third, we validate the underlying assumptions and demonstrate the pipeline’s effectiveness using a wide range of real datasets across numerous applications.

In conclusion, we consider BiPCA a significant advancement in preprocessing pipelines for high-throughput count data. By integrating biwhitening for normalization with optimal shrinkage for denoising, BiPCA provides a principled, data-driven approach that is robust across various data modalities. Its flexibility in adapting to different count distributions and its ability to preserve and enhance biological signals position BiPCA as a universal tool for biological data analysis.

## 4 Acknowledgment

Y.K. discloses support for the research of this work from NIH [R01GM131642, UM1DA051410, R33DA047037, U54AG076043, U54AG079759, P50CA121974 and U01DA053628]. D.K. discloses support for the research of this work from Gemeinnützige Hertie-Stiftung and is a member of Germany’s Excellence cluster 2064 “Machine Learning— New Perspectives for Science” (EXC 390727645).

## 5 Methods

### 5.1 BiPCA

#### 5.1.1 Marchenko-Pastur Law

We assume a low rank signal + noise model for the observed *m* × *n* data matrix *Y*, i.e.,

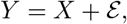

where *X*_*ij*_ = 𝔼 [*Y*_*ij*_] is a low rank (rank *r* ≪ *m*) signal matrix and ℰ is a centered noise matrix (𝔼 ℰ [_*ij*_] = 0). When the noise variables *ℰ*_*ij*_ are *homoscedastic*, i.e., *ℰ*_*ij*_ is independent and identically distributed (i.i.d.) with variance *σ* ^2^ for all *i* and *j*, the empirical spectral density of *ℰ* can be characterized using the Marchenko-Pastur (MP) distribution [35], or as known as the MP law. The probability density function of MP distribution is defined as:

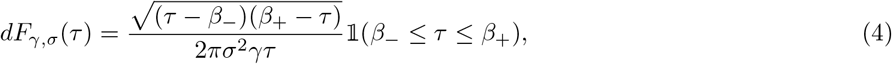

where *F* _*γ σ*_ (τ) is the cumulative density function for a MP distribution with aspect ratio γ= *m*/*n*, noise level a, and upper and lower supporting bounds 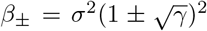. In particular, adherence to the MP by ℰ can be used to partition the data *Y* into signal and noise components: eigenvectors from the data covariance *n*^*-*1^*YY* ^*T*^ with eigenvalues larger than *β* _+_ belong to the signal components, whereas eigenvectors with eigenvalues smaller than _+_ belong to the noise components. Equivalently, singular values of *n*^*-*1^*Y* larger than 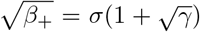 are attributable to the signal matrix *X*. whereas the singular values smaller than 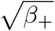 are attributable to the noise matrix ℰ. Therefore, a natural approach to estimate the rank of *X* is to count the number of singular values that are above 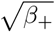.

#### 5.1.2 Biwhitening

In this work, we target data which does *not* satisfy the homoscedasticity requirements. An example is non-identical Poissons, where *Y*_*ij*_ ∼ Poisson(*X*_*ij*_) and *X*_*ij*_ is not identical across entries of *X*. In this case, it is easy to show that Var[ℰ_*ij*_] = *X*_*ij*_, hence violating the assumption of homoscedasticity. In other words, the noise matrix is *heteroscedastic*. To tailor for heteroscedastic noise in count data, we proposed *biwhitening* [36], that normalizes the data such that MP law still holds. Specifically, we normalize the rows and columns of *Y* using *whitening factors* ***û*** and 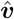:

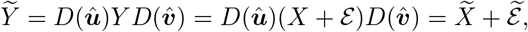

where *D*(***û***) and 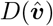 are diagonal matrix with ***û*** and 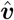 on the diagnol, 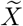and 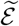 are biwhitened signal and noise matrices. Through biwhitening, the average variance is 1 in each row and column of the biwhitened noise matrix 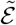, i.e.,

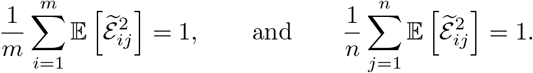

We show in [36] that after biwhitening, the spectrum of 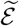 will surely converge to the MP distribution. ***û*** and 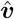are estimated through Sinkhorn-Knopp matrix scaling algorithm [36, 41]. Because non-zero diagonal scaling does not change the rank of a matrix, we can estimate the rank of 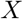 using the spectrum of 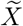 by the singular values of the biwhitened data matrix 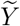 that are above 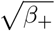 [36].

#### 5.1.3 Denoising

After biwhitening, we apply singular value shrinkage to denoise the data. Specifically, we construct the denoised signal matrix 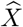 as

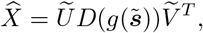

where 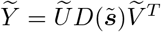 is the singular value decomposition of 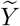 and *g* is an *optimal shrinker* ^3^. The shrinkage function *g* is derived to optimally estimate an underlying signal matrix according to a prescribed loss function. Here we consider the Frobenius shrinker *g*_*F*_, which minimizes the Frobenius loss 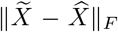 [37]. As shown in [40], the analytical form of the optimal Frobenius shrinker is given as:

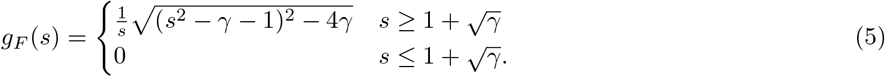

To apply the shrinker for a specific noise level a, we follow the convention in [40] to scale the singular values before shrinkage and rescale it after shrinkage. Specifically, the re-scaled shrunken singular values are computed as 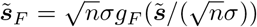, which is then used to compute the final denoised signal matrix: 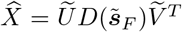. Details for estimating the noise level *σ* using the median of the empirical singular values are provided in [40]. In Section S1.1 we provide a similar procedure for estimating the noise level *σ* for a given *Y* using a quantile of its empirical spectral distribution.

#### 5.1.4 Quadratic Variance Function parameter estimation

Our approach assumes that the generating distribution of the input data *Y* has a *quadratic variance function* (QVF),

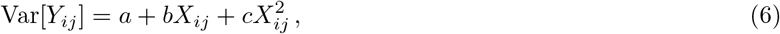

where 𝔼 [*Y*_*ij*_] = *X*_*ij*_ is the latent mean of *Y*_*ij*_ and *a, b*, and *c* are real numbers referred to as constant, linear, and quadratic variance coefficients, respectively. In [36], we propose a corresponding unbiased variance estimator 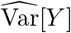 (i.e.,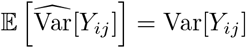 since *X* is unknown and we show that such estimator exists when *c* 6≠ −1:

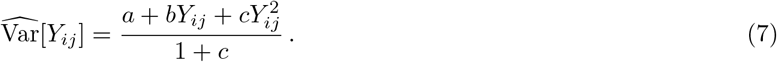

In addition, we show that one can estimate appropriate biwhitening factors ***û*** and 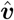 for *Y* by applying Sinkhorn-Knopp matrix scaling to the matrix of unbiased variance estimates [36]:

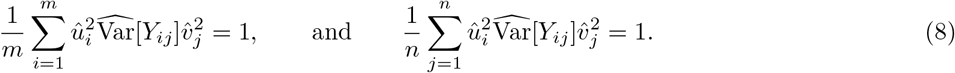

When the QVF parameters are unknown, we propose a data-driven optimization procedure to learn estimates 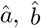 and *ĉ*. Specifically, we learn parameters that minimize a distributional distance between the biwhitened data’s empirical spectral distribution and theoretical MP. Let 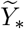 be the biwhitened data matrix when *a* _*_, *b* _*_, and *c* _*_ are used as QVF parameters. We assume that if *a* _*_, *b*_*_ and *c*_*_ are optimal, then the empirical spectral distribution of 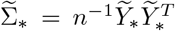 will closely follow the MP distribution with unit noise variance. We quantify this using the Kolmogorov–Smirnov (KS) distance between the empirical spectrum of 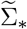 and the MP distribution,

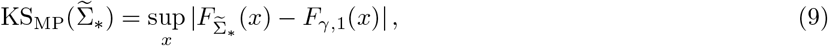

where 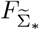 is the empirical distribution function of 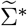 and *F*_*γ*,1_ is the cumulative distribution function for the MP distribution with aspect ratio γ= *m*/*n* and noise variance *σ*^2^ = 1. Crucially, Equation (7) and Equation (8) demonstrate that the biwhitening factors ***û*** and 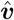 depend on the QVF parameters. Consequently, the biwhitened data matrix 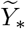 and its spectrum 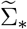 are functions of *a*_*_, *b*_*_, and *c*_*_. This allows Equation (9) to be reparameterized in terms of these parameters:

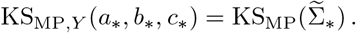

If the data *Y* is a matrix of pure noise sampled from a distribution with QVF parameters *a*_*_, *b*_*_, and *c*_*_, the empirical spectral distribution of 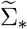 will perfectly resemble the MP distribution, and KS_MP,*Y*_ (*a* _*_, *b* _*_, *c* _*_) will approach 0. In real data, we expect there to be a rank-r latent signal matrix *X* and thus 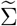 will have r eigenvalues lying above *β*_+_. In this case, the theoretically minimum achievable KS distance is *r*/*m*. Since we assume the data is low rank, this lower bound on KS resolution is typically low enough to have negligible effects on our ability to fit the data. However, to address this issue one could consider other distribution distance objectives in place of the KS.

The non-negativity of variance constrains the feasible region of each QVF parameter based on the range of *Y*_*ij*_ and the value of the other QVF parameters. This leads us to a goodness-of-fit optimization with nonlinear constraints

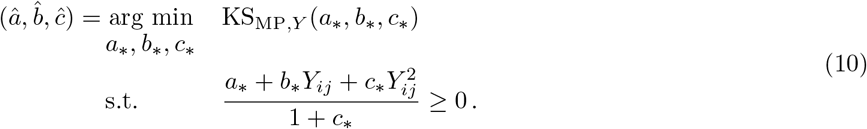

In this work, we ensure that the non-negativity constraint in Equation (10) is met by restricting our search to a widely applicable region: *a* = 0, *b* ≥ 0, and *c* ≥ 0. This reduces the complexity of our problem, as we solve

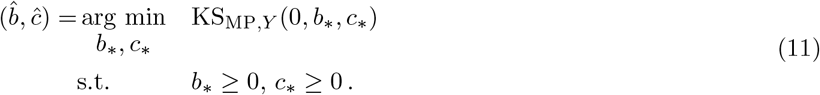

To solve Equation (11), we re-parameterize *b* and *c* such that they satisfy a convex combination with coefficient *q* up to global scaling by *σ* ^2^ (see Supplementary Methods S1.1 for details). We show that this reparametrization simplifies the objective to optimization over a single parameter *q*, for which we adopted a Chebyshev polynomials optimization procedure [64] (See Supplementary Methods S1.2 for details). After the optimization, we obtain the QVF parameters 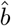 and *ĉ*, as well as the KS distance as the metric to quantify the goodness-of-fit.

Though we consider a restricted domain, this region includes many distributions we expect to fit in practice. For example, Poisson, Negative Binomial, and Gamma distributions each lie within the search space we consider. Future work could consider a larger family of distributions by solving the more complex problem of Equation (10) or adapting the parameter’s domain based on the range of the data. For example, the ensuing discussion can be immediately extended to fixed and positive estimates of *a* when *Y* is non-negative, and negative values of *a* could be used, provided that *Y* is a positive matrix and the feasible regions of *b* and *c* are suitably constrained.

#### 5.1.2 Postprocessing

After denoising, we include the following postprocessing steps to facilitate downstream analysis and interpretation.

##### Zero-thresholding

In this work, we are interested in estimating non-negative matrices. Thus, we set all negative elements in the denoised matrix 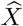 to 0 (*thresholding*). The entries of the zero-thresholded matrix 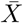 are computed as:

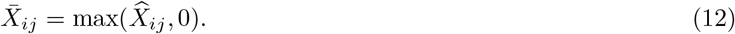

##### Library size normalization

After denoising and zero-thresholding, our postprocessing procedure includes a concluding step of library size normalization to eliminate remaining cell size factors for single cell analysis, we further include a global rescaling step to enhance interpretability.

The library normalized and globally rescaled matrix *Z* is computed as

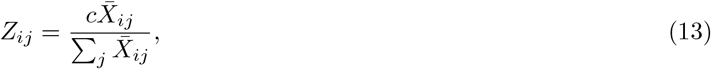

where c is a global scaling factor we set to the median library size of the original data.

##### Dimensionality reduction

We perform truncated SVD using the BiPCA estimated rank 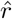 on the re-normalized data *Z* when dimensionality reduction is computationally required.

Specifically, truncated SVD estimator *Z* ^′^ is computed as:

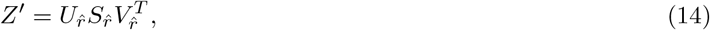

where 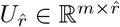 is the matrix with the first 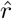 left singular vectors from 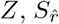 is the diagonal matrix with the top 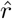 singular values from *Z*, and 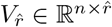 is the matrix with first 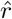 right singular vectors from *Z*. We use the loadings 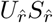 as the input to downstream analysis.

##### Dewhitening (optional)

The biwhitening step in BiPCA offers several advantages such as stabilization of the feature variance. However, one may wish to estimate the original matrix of latent means before biwhitening (i.e., in the scale of the original data) in certain applications. In this case, one can *dewhiten* the denoised data, i.e., by multiplying the rows and columns of the denoised matrix by the inverse of the whitening factors:

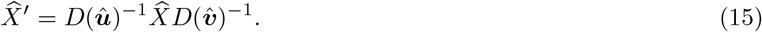

### 5.2 Data repository and processing

We collected 123 published count data sets from 40 data sources, and applied BiPCA across this collection to establish the generality of our method. The details including source of the dataset, filtering criteria are described in **Section** S2. The fitted parameters and the size of each dataset before and after filtering for each dataset is included in Supplementary table 1.

For these datasets, we filter the rows and columns of the input count matrix to remove sparse observations and features—an approach commonly adopted in many preprocessing pipelines [44, 45]. We recommend this step for several reasons. First, random matrix theory requires that each row and column of the noise matrix exhibit at least some variation in order to obtain the standard spectral behavior (MP law). Second, filtering helps our algorithms converge more reliably. Specifically, the Sinkhorn algorithm used in the biwhitening step requires a minimum density of the data for guaranteed convergence [36]. Our experiments show that mild filtering is sufficient for convergence across different modalities.

Furthermore, filtering out sparse columns and rows benefits the estimation of QVF parameters. For certain modalities that are highly sparse (e.g., scATAC-seq), we found that more filtering would reduce the variance of the estimates. However, there is a trade-off, as more aggressive filtering may remove informative features. For instance, in the experiment illustrated in Fig. 5c, we applied lighter filtering to the scATAC-seq data to preserve enough informative features for accurate cell-type classification. Even so, we observed stable parameter estimations with mild filtering across most datasets (see Fig. 2d). We recommend that users carefully consider their sparsity filtering criteria as a balance between algorithmic stability and signal extraction.

### 5.3 Comparison methods

For scRNA-seq datasets, we compared BiPCA with five normalization methods: log-normalization (log1p), log-normalization with gene-wise standardization (log1p+z), Pearson residuals [28] (implemented in Scanpy [45]), Sanity [29] (https://github.com/jmbreda/Sanity), ALRA [46] (https://github.com/milescsmith/pyalra). For scATAC-seq datasets, we compared BiPCA with three normalization methods: log-normalization (log1p), log-normalization with peak-wise standardization (log1p+z), and Term Frequency - Inverse Document Frequency (TF-IDF) (implemented in muon [47]).

In tasks that involved low-rank approximations, we selected the approximation rank using standard practices for each method unless otherwise noted. For log1p, log1p+z, Pearson, Sanity, and TF-IDF, the rank was set to 50. For ALRA, the rank was set based on the proposed rank estimation procedure [46]. For BiPCA, the rank was set according to the pipeline’s estimate.

### 5.4 Simulations

#### 5.4.1 Simulation of prescribed rank and QVF parameters

##### Simulation of Poisson-sampled matrices of prescribed rank

To evaluate the accuracy of rank estimation, we simulated 5000 × 5000 Poisson matrices with a grid of underlying ranks {1, 2, 4, …, 2^*p*^}, where *p* = 0, 1,…, 6. For a matrix of rank r, we first sampled library size factors *N*_*i*_ for each row using a log-normal distribution:

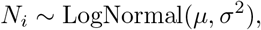

where µ = log(1000) and *σ* ^2^ = 0.01. Using these library sizes, we generated a signal matrix *U* ℝ ^5000× *r*^ in which each row vector was sampled from a multinomial distribution:

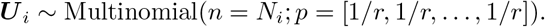

Next, we sampled a coefficient matrix *V* ^*T*^ ∈ ℝ ^5000× *r*^ where each 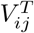 was uniformly sampled at random from a logarithmic spacing of 5000r values between 1 × 10^*-*4^ to 5 × 10^*-*2^. The raw signal matrix is denoted as *X*^′^ = *UV* ^*T*^, and we constructed a final ground truth signal matrix X with entrywise mean λ by rescaling *X*^′^:

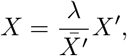

where 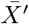 is the mean of *X*^′^. The simulated data matrix *Y* was then sampled according to:

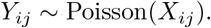

In this experiment, we set λ to 20, which we found maintained a reasonable signal-to-noise ratio in the data. We found that as rank increases for a fixed mean, the total signal in each component becomes diluted, making rank estimation more challenging as SNR degrades. Larger λ values would be more appropriate in such regimes.

##### Simulation of rank 1 random matrices with prescribed QVF parameters

To evaluate QVF parameter estimation performance, we similarly simulated 5000 × 5000 matrices but used different ground truth *b* and *c* and fixed *r* = 1. The signal matrix *X* was generated as described above. However, the data matrix *Y* was generated using a scaled negative binomial distribution:

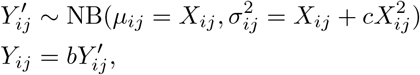

where µ and *σ*^2^ denote the mean and variance of the negative binomial distribution. When estimating parameter *b*, we fixed *c* = 1 × 10^*-*6^ and varied *b* over {2^*-*7^, 2^*-*6^,…, 2^6^}. When estimating parameter *c*, we fixed *b* = 1 and varied c over {2^*-*7^, 2^*-*6^,…, 2^6^}.

#### 5.4.2 Simulation of rank-5 data

We simulate a rank-5 count matrix with 3000 observations and 1000 features. First, we generate 5 vectors *e*_1_, …, *e*_5_ ∈ ℝ^200^ where e_*i*_(j) ∼ Exp(1), from which we generate 5 linearly independent signal vectors v_1_, …, v_5_ 2 ℝ^1000^ where

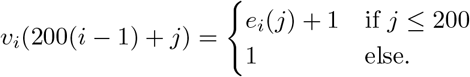

Let *V* be the matrix of signal vectors, i.e, *V* = [*v*_1_ *v*_2_ *v*_5_]. The coefficients *β*∈*R*^3000v5^ are sampled from multinomial(0.2, 0.2, 0.2, 0.2, 0.2) that indicate the ground truth cluster labels. The count matrix *X* is generated from a Poisson distribution where *X* ∼ Poisson((*µ βV*)) where µ is a scaling factor and Signal-to-Noise Ratio (SNR) is defined as 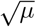.We vary the scaling factor *µ* = {1, 2, 4, ‥, 2^*p*^} where *p* ∈ {0‥9} to change SNR and compare 3 rank approximation approaches: rank-30 approximation, rank-50 approximation, BiPCA adaptive rank approximation.

Silhouette scores are computed with respect to the ground truth cluster labels for each rank approximation approach at each SNR regime. µ = 2 is used for comparing the t-SNEs between rank-50 approximation and BiPCA rank approximation.

### 5.5 Analysis of proper rank estimation

#### 5.5.1 Data processing

##### Zheng2017 PBMCs

From the preprocessed *Zheng2017* data detailed in Section S2, we filter out cells that have less than 100 genes expressed and more than 10% mitochondrial genes. We also filter out genes that have less than 100 cells expressed. The filtered data includes 94, 572 cells and 12, 791 genes.

##### PFC 10x Multiome

For each processed sample detailed in Section S2, we run leiden clustering with resolution = 3 using `scanpy` [45] on the BiPCA low rank approximated data with the number of PCs specified by BiPCA, and annotate the Oligodendrocyte Precursor Cells (OPC) clusters using marker gene PDGFRA.

#### 5.5.2 Identification of the cell subtypes and validation

To identify cell subtypes associated with weaker signal components, we first run leiden clustering with resolution = 3 on the BiPCA low rank approximated data with signal components that are out of the top 50. We identify a cluster of B cells within the CD19+ B cell population in the Zheng2017 PBMC data that have unique TCL1B expression. Therefore, we term this population TCL1B+ B cells. Similarly, we identify a cluster of OPCs within the OPCs in the human PFC data that have unique SEMA3E expression, for which we termed SEMA3E+ OPCs.

To validate the identified cell type sub-clusters (TCL1B+ B cells and SEMA3E+ OPCs). We compute the relative energy (RE) for each sub-cluster relative to the parent population (i.e., all B cells or all OPCs) across singular vectors to measure how well each singular vector differentiates each sub-cluster from the whole population.

Specifically, the energy for each component i is defined as the l_2_ norm of the singular vector *U*_*i*_ scaled by its singular value *S*_*i*_, then normalized by the total norm.

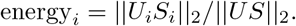

The relative energy for the cells in sub-cluster (indexed as sub) relative to the parent cell population (indexed as parent) is defined as:

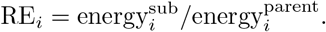

We performed a Mann-Whitney U test comparing each sub-cluster (e.g., TCL1B+ B cells) to the other cells in the parent population (e.g., the other *B* cells) to identify marker genes associated with each sub-cluster. We select the strongest marker genes with logFC > 1 and adjusted p-values < 0.01 (Bonferroni correction).

### 5.6 Quantification of cell neighborhood preservation

For this task, we focus on three sets of datasets: (1) CITE-seq (RNA): The RNA modality of 14 CITE-seq datasets including *Stoekius2017* [53], *Stuart2019* [54], and 12 batches from *Luecken2021CITE* [55], (2) Multiome (RNA): The RNA modality of 13 10x Multiome datasets (13 batches from *Luecken2021Multiome* [55]), and (3) Multiome (ATAC): The ATAC modality of the same 13 10x Multiome datasets. The processing steps of each dataset are described in detail in Section S2.

For each dataset, we first apply baseline normalization methods to the filtered count matrices, then we apply low rank approximation from the normalized data of each method to reduce the dimensions, with *r* specified as in **Section** 5.3 for each method. We use the cell types as the cluster labels and compute silhouette scores on the low dimensional space [56, 65]. For each cell i, we first compute the mean intra-cluster distance between cell i and other cells in its cluster, defined as

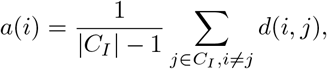

where *C*_*I*_ is the set of cells of the cluster cell i is in, |*C*_*I*_| is the number of cells in that cluster, and *d*(*i, j*) is the euclidean distance between cell *i* and *j* on the low dimensional space. Next, we compute the mean distance between cell *i* and the cells from the nearest other cluster. for each cell *i*:

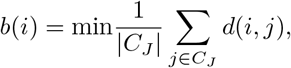

where *J* ≠ *I*. The silhouette score *s*(*i*) is defined as

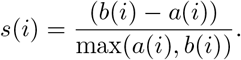

We used the mean silhouette scores for all cells as the final metric. A method will have higher scores if it has small intra-cluster distances and large inter-cluster distances.

In addition, we computed KNN accuracy on the low dimensional space [56, 65] to measure how well neighborhoods were preserved. We fixed *k* = 10 and predicted cell labels from the mode of their *k* nearest neighbors. Balanced accuracy over labels is For computing both silhouette scores and KNN accuracy, we subsampled 80% of the data and repeated each experiment 10 times. We reported the mean and standard deviations in the final results.

### 5.7 Marker gene analysis

#### 5.7.1 Data processing

From the preprocessed *Zheng2017* data detailed in Section S2, we coarse-grained the cluster labels into four major cell types: CD19+ B cells, CD4+ T cells, CD8+ T cells, and CD56+ NK cells and removed other cell types. We then filter out cells that have less than 100 genes expressed and more than 10% mitochondrial genes. We also filter out genes that have less than 100 cells expressed. The filtered data includes 82, 746 cells and 11, 539 genes.

To comprehensively benchmark the coherence of marker gene expressions with the corresponding cell types, we extracted a set of 101 marker gene annotations from Human Protein Atlas (HPA) [50]. First, we filtered out genes that are not detected in blood or immune cells and genes that have low specificity, and keep only genes that are enriched in CD19+ B cells, CD56+ NK cells, CD4+ and CD8+ T cells. We obtained in total 101 marker gene annotations, with 65 B cell markers, 21 NK cell markers, and 15 T cell markers, respectively.

#### 5.7.2 Marker density and area under the receiver operating characteristic curve (AUROC)

We plotted the kernel density for CD40, CD4, CD8A, and CD56, which are canonical marker genes for CD19+ B cells, CD4+ T cells, CD8+ T cells, and CD56+ NK cells, respectively. The expression values for each gene and transformation pair were feature scaled into the range [0, 1]. Two gene-wise kernel density estimates were computed over the scaled data: one over all the cells in the positive cluster (i.e., the cell type that corresponds to that gene), and a second over all the cells in the complementary negative cluster. Density estimation was performed using the `gaussian kde` function implemented by the Python package `scipy`. For visualization, the kernel density estimates were evaluated over a 1000 point grid on [0, 1] and the resulting densities were independently feature scaled to produce curves of similar height.

We compute the area under the receiver operating characteristic curve (AUROC) for each gene-transformation pair using the `roc auc score` function from scikit-learn [65]. Specifically, `roc auc score` accepts two inputs: x, a vector of transformed gene expression values across all cells, and y, a binary label vector indicating whether each cell belongs to the positive or negative cluster.

### 5.8 Quantification of batch effects

We analyzed a 10x Chromium V3 single-nucleus RNA-seq data (*Kluger2024OUD*) of human insular cortex (INS) from the SCORCH consortium [51], where we identified batch effects across 4 technical replicates. The data processing steps are detailed in Section S2. For the processed data, we run leiden clustering using `scanpy` [45] on the BiPCA low rank approximated data with the number of PCs specified by BiPCA, and annotate the Astrocytes clusters using marker gene AQP4. We ran log1p, log1p+z, Pearson, Sanity, ALRA, and BiPCA on this data and quantified the batch effect with three metrics: number of differentially expressed genes, Laplacian score, and Affine Grassmann distance [58, 59] between batches.

#### 5.8.1 Differential gene expression analysis

For each gene, we performed Mann-Whitney U test between replicate 1 and other replicates. We select differentially expressed genes with adjusted p-values < 0.01 (Bonferroni correction).

#### 5.8.2 Laplacian score

Let *x*_*i*_ and *x*_*j*_ denote the expression profile of the *i* th and the *j* th cell, the affinity between *x*_*i*_ and *x*_*j*_ in the affinity matrix *K* with a Gaussian kernel is defined as:

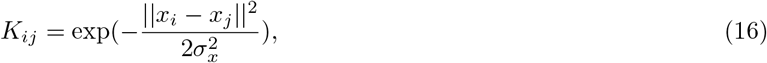

where 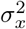 is the kernel bandwidth of the local affinity. And graph Laplacian is defined as:

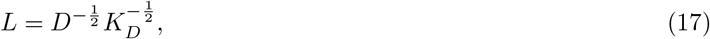

where 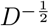 is a diagonal matrix of the row sums of *K*. An important property of graph Laplacian is that its top eigenvectors that correspond to large eigenvalues reflect the underlying geometry of the data, which is utilized by Laplacian score [57] to measure the smoothness of features in the data. Specifically, the Laplacian score of feature vector *f* with respect to *L*_*x*_ is defined as 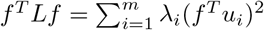 where 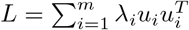 is the eigen-decomposition of *L*. If feature *f* is low frequency, i.e., it varies slowly along the underlying structure, the corresponding Laplacian score will be high. Otherwise it will have a low score.

To quantify how well replicate 1 mix with the other replicates in the astrocyte cluster, we define indicator vector f ∈ℝ^*m*^ where 1 indicates that cell belongs to replicate 1 and 0 otherwise. For each method, we construct the graph Laplacian *L* and and compute Laplacian score *S*:

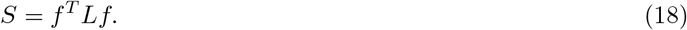

The smaller the score metric *S*, the better the replicates are mixed together in the cluster for that method.

#### 5.8.3 Affine Grassman distance

##### Affine PCA Coordinates

Let *X* ∈ ℝ^*m*×*n*^ denote the normalized data matrix after each transformation. We first compute the mean vector 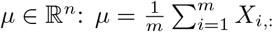,and perform SVD on the mean-centered matrix *X*_*c*_:

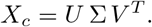

Next, we project µ onto the orthogonal complement of *V*_*r*_ (first r right singular vectors of *V*):

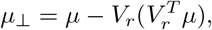

and define the scaling factor γ as 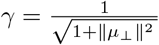.Together, 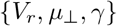,characterizes the affine subspace µ+span(*V*_*r*_).

##### Embedding into the Stiefel Manifold

Next, we embed the affine subspace µ+span(*V*_*r*_) into an (*n*+ 1) × (*r* + 1) matrix *Y* via

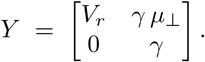

The columns of *Y* are orthonormal in a lifted space ℝ^*n*+1^. Hence, *Y* ∈ *St* (*r* + 1, *n* + 1), where *St*(*k, n*) denotes the Stiefel manifold of all orthonormal *n* v *k* matrices.

##### Principal Angles between Two Subspaces and Affine Grassmann Distance

Since we want to measure the differences between replicate 1 and cells from the other replicates, we split *X* into two subsets - *X*^(1)^ (all cells from replicate 1) and *X*^(2)^ (all cells from other replicates) and then compute the affine subspace for each, respectively. The corresponding embeddings for each subset are as follows:

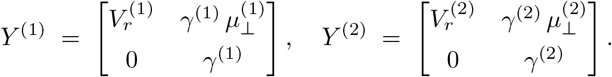

We then compute an SVD of

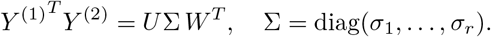

The principal angles θ_*i*_ between the two affine subspaces are given by

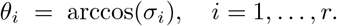

Finally, the affine grassmann distance between the two affine subspaces is computed as

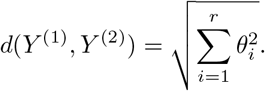

A distance of zero indicates that *Y* ^(1)^ and *Y* ^(2)^ span the same affine subspace, whereas larger values reflect greater differences in both the orientation and the offset of these subspaces. In other words, if replicate 1 and other replicates are mixed well, their corresponding distance will be small, indicating small batch effect between them after data normalization.

### 5.9 Quantification of Mean-Variance relationships

We analyzed one human PBMC scRNA-seq dataset, *10×2016PBMC*, from 10x Geomics [66]. We filter the data (detailed in Section S2) and then apply the aforementioned normalization methods to transform the data.

#### 5.9.1 Mean-variance relationship after different transformations

We first examined the mean-variance relationship on the transformations that do not involve low rank approximation: log1p, Sanity, Pearson, and Biwhitening. For each gene *g*, its mean expression across all cells is measured on the raw data 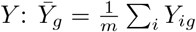 whereas its variance across cells is computed after each transformation *T* : Var(*T* (*Y*)_*g*_) where Var() is the empirical variance function. Next, we focused on all transformations coupled with low rank approximation: log1p, log1p+z, Pearson, ALRA, Sanity, and BiPCA, with rank r specified as described in Section 5.3. After low rank approximations, we followed the same procedure to compute the mean and variance and examined their relationship.

#### 5.9.2 Assessing the stability of gene variances

To measure the stability of the gene variances after each transformation, we subsampled 5000 cells and re-applied each transformation *T* to the subsampled data along with low rank approximation. For each gene g, we compare its relative variance on the full data: 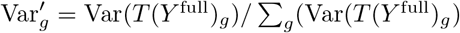,to its relative variance on the subsampled data: 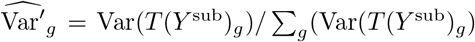.Lastly, we computed the spearman correlation coefficient *r*_*s*_ between Var^*′*^ and 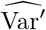 across all genes as the final metric.

## Footnotes

1 For this exposition, we assume *m ≤ n*; we can always transpose the data in practice.

2 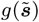 is the vector formed by evaluating the scalar function *g* : ℝ *7!* ℝ on every element of the vector 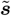.

3 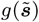 is the vector formed by evaluating the scalar function *g* : ℝ ⟼ ℝ on every element of the vector 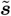.

## References

[1] Andrea Riba, Attila Oravecz, Matej Durik, Sara Jimenez, Violaine Alunni, Marie Cerciat, Matthieu Jung, Celine Keime, William M Keyes, and Nacho Molina. Cell cycle gene regulation dynamics revealed by rna velocity and deep-learning. Nature communications, 13(1): 2865, 2022.

[2] Emily Stephenson, Gary Reynolds, Rachel A Botting, Fernando J Calero-Nieto, Michael D Morgan, Zewen Kelvin Tuong, Karsten Bach, Waradon Sungnak, Kaylee B Worlock, Masahiro Yoshida, et al. Single-cell multi-omics analysis of the immune response in covid-19. Nature medicine, 27(5):904–916, 2021.

[3] Marta Byrska-Bishop, Uday S Evani, Xuefang Zhao, Anna O Basile, Haley J Abel, Allison A Regier, Andre Corvelo, Wayne E Clarke, Rajeeva Musunuri, Kshithija Nagulapalli, et al. High-coverage whole-genome sequencing of the expanded 1000 genomes project cohort including 602 trios. Cell, 185(18):3426–3440, 2022.

[4] Mayo Blegen Ashley L. 18 Wirkus Samantha J. 18 Wagner Victoria A. 18 Meyer Jeffrey G. 18 Cicek Mine S. 10 18 Biobank and All of Us Research Demonstration Project Teams Choi Seung Hoan 14 http://orcid.org/0000-0002-0322-8970 Wang Xin 14 http://orcid.org/00000001-6042-4487 Rosenthal Elisabeth A. 15. Genomic data in the all of us research program. Nature, 627(8003):340–346, 2024.

[5] Takashi Nagano, Yaniv Lubling, Tim J Stevens, Stefan Schoenfelder, Eitan Yaffe, Wendy Dean, Ernest D Laue, Amos Tanay, and Peter Fraser. Single-cell hi-c reveals cell-to-cell variability in chromosome structure. Nature, 502(7469):59–64, 2013.

[6] Darren A Cusanovich, Andrew J Hill, Delasa Aghamirzaie, Riza M Daza, Hannah A Pliner, Joel B Berletch, Galina N Filippova, Xingfan Huang, Lena Christiansen, William S DeWitt, et al. A single-cell atlas of in vivo mammalian chromatin accessibility. Cell, 174(5):1309–1324, 2018.

[7] Dragomirka Jovic, Xue Liang, Hua Zeng, Lin Lin, Fengping Xu, and Yonglun Luo. Single-cell rna sequencing technologies and applications: A brief overview. Clinical and translational medicine, 12(3):e694, 2022.

[8] Lambda Moses and Lior Pachter. Museum of spatial transcriptomics. Nature methods, 19(5):534–546, 2022.

[9] Alev Baysoy, Zhiliang Bai, Rahul Satija, and Rong Fan. The technological landscape and applications of single-cell multi-omics. Nature Reviews Molecular Cell Biology, 24(10):695–713, 2023.

[10] Katy Vandereyken, Alejandro Sifrim, Bernard Thienpont, and Thierry Voet. Methods and applications for single-cell and spatial multi-omics. Nature Reviews Genetics, 24(8):494–515, 2023.

[11] Oliver Stegle, Sarah A Teichmann, and John C Marioni. Computational and analytical challenges in single-cell transcriptomics. Nature Reviews Genetics, 16(3):133–145, 2015.

[12] David Lahnemann, Johannes Koster, Ewa Szczurek, Davis J McCarthy, Stephanie C Hicks, Mark D Robinson, Catalina A Vallejos, Kieran R Campbell, Niko Beerenwinkel, Ahmed Mahfouz, et al. Eleven grand challenges in single-cell data science. Genome biology, 21:1–35, 2020.

[13] Laura D Martens, David S Fischer, Vicente A Yepez, Fabian J Theis, and Julien Gagneur. Modeling fragment counts improves single-cell atac-seq analysis. Nature Methods, 21(1):28–31, 2024.

[14] Malte D Luecken and Fabian J Theis. Current best practices in single-cell rna-seq analysis: a tutorial. Molecular systems biology, 15(6):e8746, 2019.

[15] Lukas Heumos, Anna C Schaar, Christopher Lance, Anastasia Litinetskaya, Felix Drost, Luke Zappia, Malte D Lucken, Daniel C Strobl, Juan Henao, Fabiola Curion, et al. Best practices for single-cell analysis across modalities. Nature Reviews Genetics, 24(8):550–572, 2023.

[16] Constantin Ahlmann-Eltze and Wolfgang Huber. Comparison of transformations for single-cell rna-seq data. Nature Methods, 20(5):665–672, 2023.

[17] Lucrezia Patruno, Davide Maspero, Francesco Craighero, Fabrizio Angaroni, Marco Antoniotti, and Alex Graudenzi. A review of computational strategies for denoising and imputation of single-cell transcriptomic data. Briefings in bioinformatics, 22(4):bbaa222, 2021.

[18] Lijia Yu, Yue Cao, Jean YH Yang, and Pengyi Yang. Benchmarking clustering algorithms on estimating the number of cell types from single-cell rna-sequencing data. Genome biology, 23(1): 49, 2022.

[19] Hoa Thi Nhu Tran, Kok Siong Ang, Marion Chevrier, Xiaomeng Zhang, Nicole Yee Shin Lee, Michelle Goh, and Jinmiao Chen. A benchmark of batch-effect correction methods for single-cell rna sequencing data. Genome biology, 21:1–32, 2020.

[20] Wenpin Hou, Zhicheng Ji, Hongkai Ji, and Stephanie C Hicks. A systematic evaluation of single-cell rna-sequencing imputation methods. Genome biology, 21:1–30, 2020.

[21] Joseph M Rich, Lambda Moses, Petur Helgi Einarsson, Kayla Jackson, Laura Luebbert, A Sina Booeshaghi, Sindri Antonsson, Delaney K Sullivan, Nicolas Bray, Pall Melsted, et al. The impact of package selection and versioning on single-cell rna-seq analysis. bioRxiv, 2024.

[22] Romain Lopez, Jeffrey Regier, Michael B Cole, Michael I Jordan, and Nir Yosef. Deep generative modeling for single-cell transcriptomics. Nature methods, 15(12):1053–1058, 2018.

[23] Davide Risso, Fanny Perraudeau, Svetlana Gribkova, Sandrine Dudoit, and Jean-Philippe Vert. A general and flexible method for signal extraction from single-cell rna-seq data. Nature communications, 9(1): 284, 2018.

[24] Gokcen Eraslan, Lukas M Simon, Maria Mircea, Nikola S Mueller, and Fabian J Theis. Single-cell rna-seq denoising using a deep count autoencoder. Nature communications, 10(1): 390, 2019.

[25] Valentine Svensson. Droplet scrna-seq is not zero-inflated. Nature Biotechnology, 38(2):147–150, 2020.

[26] F William Townes, Stephanie C Hicks, Martin J Aryee, and Rafael A Irizarry. Feature selection and dimension reduction for single-cell rna-seq based on a multinomial model. Genome biology, 20(1):1–16, 2019.

[27] Christoph Hafemeister and Rahul Satija. Normalization and variance stabilization of single-cell rna-seq data using regularized negative binomial regression. Genome biology, 20(1): 296, 2019.

[28] Jan Lause, Philipp Berens, and Dmitry Kobak. Analytic pearson residuals for normalization of single-cell rna-seq umi data. Genome biology, 22(1):1–20, 2021.

[29] Jeremie Breda, Mihaela Zavolan, and Erik van Nimwegen. Bayesian inference of gene expression states from single-cell rna-seq data. Nature Biotechnology, 39(8):1008–1016, 2021.

[30] Luis Aparicio, Mykola Bordyuh, Andrew J Blumberg, and Raul Rabadan. A random matrix theory approach to denoise single-cell data. Patterns, 1(3), 2020.

[31] Mor Nitzan and Michael P Brenner. Revealing lineage-related signals in single-cell gene expression using random matrix theory. Proceedings of the National Academy of Sciences, 118(11):e1913931118, 2021.

[32] Maria Mircea, Mazene Hochane, Xueying Fan, Susana M Chuva de Sousa Lopes, Diego Garlaschelli, and Stefan Semrau. Phiclust: a clusterability measure for single-cell transcriptomics reveals phenotypic subpopulations. Genome Biology, 23:1–24, 2022.

[33] Hyun Kim, Won Chang, Seok Joo Chae, Jong-Eun Park, Minseok Seo, and Jae Kyoung Kim. sclens: data-driven signal detection for unbiased scrna-seq data analysis. Nature Communications, 15(1): 3575, 2024.

[34] Sivan Leviyang. Analysis of a single cell rna-seq workflow by random matrix theory methods. Bulletin of Mathematical Biology, 87(1): 4, 2025.

[35] Vladimir A Marcenko and Leonid Andreevich Pastur. Distribution of eigenvalues for some sets of random matrices. Mathematics of the USSR-Sbornik, 1(4): 457, 1967.

[36] Boris Landa, Thomas TCK Zhang, and Yuval Kluger. Biwhitening reveals the rank of a count matrix. SIAM journal on mathematics of data science, 4(4):1420–1446, 2022.

[37] Andrey A Shabalin and Andrew B Nobel. Reconstruction of a low-rank matrix in the presence of gaussian noise. Journal of Multivariate Analysis, 118:67–76, 2013.

[38] Raj Rao Nadakuditi. Optshrink: An algorithm for improved low-rank signal matrix denoising by optimal, data-driven singular value shrinkage. IEEE Transactions on Information Theory, 60(5):3002–3018, 2014.

[39] David L Donoho and Matan Gavish. The optimal hard threshold for singular values is 4/sqrt (3). arXiv preprint arXiv:1305.5870, 2013.

[40] Matan Gavish and David L Donoho. Optimal shrinkage of singular values. IEEE Transactions on Information Theory, 63(4):2137–2152, 2017.

[41] Richard Sinkhorn and Paul Knopp. Concerning nonnegative matrices and doubly stochastic matrices. Pacific Journal of Mathematics, 21(2):343–348, 1967.

[42] Carl N Morris. Natural exponential families with quadratic variance functions. The Annals of Statistics, pages 65–80, 1982.

[43] Jason D Buenrostro, Beijing Wu, Ulrike M Litzenburger, Dave Ruff, Michael L Gonzales, Michael P Snyder, Howard Y Chang, and William J Greenleaf. Single-cell chromatin accessibility reveals principles of regulatory variation. Nature, 523(7561):486–490, 2015.

[44] Yuhan Hao, Tim Stuart, Madeline H Kowalski, Saket Choudhary, Paul Hoffman, Austin Hartman, Avi Srivastava, Gesmira Molla, Shaista Madad, Carlos Fernandez-Granda, et al. Dictionary learning for integrative, multimodal and scalable single-cell analysis. Nature biotechnology, 42(2):293–304, 2024.

[45] F Alexander Wolf, Philipp Angerer, and Fabian J Theis. Scanpy: large-scale single-cell gene expression data analysis. Genome biology, 19:1–5, 2018.

[46] George C Linderman, Jun Zhao, and Yuval Kluger. Zero-preserving imputation of scrna-seq data using low-rank approximation. BioRxiv, page 397588, 2018.

[47] Danila Bredikhin, Ilia Kats, and Oliver Stegle. Muon: multimodal omics analysis framework. Genome biology, 23(1): 42, 2022.

[48] NanoString Technologies. Cosmx human frontal cortex ffpe dataset. Available at https://nanostring.com/products/cosmx-spatial-molecular-imager/ffpe-dataset/human-frontal-cortex-ffpe-dataset/, September 2023. [Online; accessed 30-July-2024].

[49] Grace XY Zheng, Jessica M Terry, Phillip Belgrader, Paul Ryvkin, Zachary W Bent, Ryan Wilson, Solongo B Ziraldo, Tobias D Wheeler, Geoff P McDermott, Junjie Zhu, et al. Massively parallel digital transcriptional profiling of single cells. Nature communications, 8(1):1–12, 2017.

[50] The Human Protein Atlas Version 21. Human Protein Atlas. Available at https://www.proteinatlas.org. x[Online; accessed 5-January-2024].

[51] Seth A Ament, Rianne R Campbell, Mary Kay Lobo, Joseph P Receveur, Kriti Agrawal, Alejandra Borjabad, Siddappa N Byrareddy, Linda Chang, Declan Clarke, Prashant Emani, et al. The single-cell opioid responses in the context of hiv (scorch) consortium. Molecular Psychiatry, pages 1–12, 2024.

[52] Kevin R Moon, Jay S Stanley III, Daniel Burkhardt, David van Dijk, Guy Wolf, and Smita Krishnaswamy. Manifold learning-based methods for analyzing single-cell rna-sequencing data. Current Opinion in Systems Biology, 7:36–46, 2018.

[53] Marlon Stoeckius, Christoph Hafemeister, William Stephenson, Brian Houck-Loomis, Pratip K Chattopadhyay, Harold Swerdlow, Rahul Satija, and Peter Smibert. Simultaneous epitope and transcriptome measurement in single cells. Nature methods, 14(9):865–868, 2017.

[54] Tim Stuart, Andrew Butler, Paul Hoffman, Christoph Hafemeister, Efthymia Papalexi, William M Mauck, Yuhan Hao, Marlon Stoeckius, Peter Smibert, and Rahul Satija. Comprehensive integration of single-cell data. Cell, 177(7):1888–1902, 2019.

[55] Malte D Luecken, Daniel Bernard Burkhardt, Robrecht Cannoodt, Christopher Lance, Aditi Agrawal, Hananeh Aliee, Ann T Chen, Louise Deconinck, Angela M Detweiler, Alejandro A Granados, et al. A sandbox for prediction and integration of dna, rna, and proteins in single cells. In Thirty-fifth conference on neural information processing systems datasets and benchmarks track (Round 2), 2021.

[56] Jan Lause, Philipp Berens, and Dmitry Kobak. The art of seeing the elephant in the room: 2d embeddings of single-cell data do make sense. bioRxiv, 2024.

[57] Xiaofei He, Deng Cai, and Partha Niyogi. Laplacian score for feature selection. Advances in neural information processing systems, 18, 2005.

[58] Lek-Heng Lim, Ken Sze-Wai Wong, and Ke Ye. The grassmannian of affine subspaces. Foundations of Computational Mathematics, 21:537–574, 2021.

[59] Krishan Sharma and Renu Rameshan. Image set classification using a distance-based kernel over affine grassmann manifold. IEEE transactions on neural networks and learning systems, 32(3):1082–1095, 2020.

[60] Tim Stuart, Avi Srivastava, Shaista Madad, Caleb A Lareau, and Rahul Satija. Single-cell chromatin state analysis with signac. Nature methods, 18(11):1333–1341, 2021.

[61] David Hong, Laura Balzano, and Jeffrey A Fessler. Asymptotic performance of pca for high-dimensional heteroscedastic data. Journal of multivariate analysis, 167:435–452, 2018.

[62] Boris Landa and Yuval Kluger. The dyson equalizer: Adaptive noise stabilization for low-rank signal detection and recovery. Information and Inference: A Journal of the IMA, 14(1):iaae036, 2025.

[63] Abhishek Sarkar and Matthew Stephens. Separating measurement and expression models clarifies confusion in single-cell RNA sequencing analysis. Nature Genetics, 53(6):770–777, 2021.

[64] John C Mason and David C Handscomb. Chebyshev polynomials. Chapman and Hall/CRC, 2002.

[65] F. Pedregosa, G. Varoquaux, A. Gramfort, V. Michel, B. Thirion, O. Grisel, M. Blondel, P. Prettenhofer, R. Weiss, V. Dubourg, J. Vanderplas, A. Passos, D. Cournapeau, M. Brucher, M. Perrot, and E. Duchesnay. Scikit-learn: Machine learning in Python. Journal of Machine Learning Research, 12:2825–2830, 2011.

[66] 10X Genomics. 33k pbmcs from a healthy donor. Available at https://www.10xgenomics.com/resources/datasets/33-k-pbm-cs-from-a-healthy-donor-1-standard-1-1-0, September 2016. [Online; accessed 17-April-2023].

[67] Michael Hagemann-Jensen, Christoph Ziegenhain, Ping Chen, Daniel Ramskold, Gert-Jan Hendriks, Anton JM Larsson, Omid R Faridani, and Rickard Sandberg. Single-cell rna counting at allele and isoform resolution using smart-seq3. Nature biotechnology, 38(6):708–714, 2020.

[68] Michael Hagemann-Jensen, Christoph Ziegenhain, and Rickard Sandberg. Scalable single-cell rna sequencing from full transcripts with smart-seq3xpress. Nature Biotechnology, 40(10):1452–1457, 2022.

[69] 10X Genomics. 9k brain cells from an e18 mouse. Available at https://www.10xgenomics.com/resources/datasets/9-k-brain-cells-from-an-e-18-mouse-2-standard-1-3-0, February 2017. [Online; accessed 17-April-2023].

[70] 10X Genomics. 8k pbmcs from a healthy donor. Available at https://www.10xgenomics.com/resources/datasets/8-k-pbm-cs-from-a-healthy-donor-2-standard-1-3-0, February 2017. [Online; accessed 17-April-2023].

[71] 10X Genomics. 20k 1:1 mixture of human hek293t and mouse nih3t3 cells, 5’ ht v2.0. Available at url https://www.10xgenomics.com/resources/datasets/20-k-1-1-mixture-of-human-hek-293-t-and-mouse-nih-3-t-3-cells-5-ht-v-2-0-2-high-6-1-0, August 2021. [Online; accessed 30-May-2023].

[72] 10X Genomics. 10k brain cells from an e18 mouse (v3 chemistry). Available at https://www.10xgenomics.com/resources/datasets/10-k-brain-cells-from-an-e-18-mouse-v-3-chemistry-3-standard-3-0-0, November 2018. [Online; accessed 17-April-2023].

[73] 10X Genomics. 10k pbmcs from a healthy donor (v3 chemistry). Available at https://www.10xgenomics.com/resources/datasets/10-k-pbm-cs-from-a-healthy-donor-v-3-chemistry-3-standard-3-0-0, Novem-ber 2018. [Online; accessed 17-April-2023].

[74] Pierre-Luc Germain, Aaron Lun, Carlos Garcia Meixide, Will Macnair, and Mark D Robinson. Doublet identification in single-cell sequencing data using scdblfinder. F1000Research, 10, 2021.

[75] 10X Genomics. 10k mouse e18 combined cortex, hippocampus and subventricular zone cells, dual in-dexed. Available at https://www.10xgenomics.com/resources/datasets/10-k-mouse-e-18-combined-cortex-hippocampus-and-subventricular-zone-cells-dual-indexed-3-1-standard-4-0-0, July 2020. [Online; accessed 17-April-2023].

[76] 10X Genomics. 20k 1:1 mixture of human hek293t and mouse nih3t3 cells, 3’ ht v3.1. Available at url https://www.10xgenomics.com/resources/datasets/20-k-1-1-mixture-of-human-hek-293-t-and-mouse-nih-3-t-3-cells-3-ht-v-3-1-3-1-high-6-1-0, August 2021. [Online; accessed 30-May-2023].

[77] 10X Genomics. 20k human pbmcs, 3’ ht v3.1, chromium x. Available at https://www.10xgenomics.com/resources/datasets/20-k-human-pbm-cs-3-ht-v-3-1-chromium-x-3-1-high-6-1-0, August 2021. [On-line; accessed 17-April-2023].

[78] Rahul Satija, Paul Hoffman, and Andrew Butler. SeuratData: Install and Manage Seurat Datasets, 2019. http://www.satijalab.org/seurat, https://github.com/satijalab/seurat-data.

[79] 10x Genomics. Human Breast Cancer (block a sections 1 and 2). Available at https://www.10xgenomics.com/resources/datasets/human-breast-cancer-block-a-section-1-1-standard-1-1-0 and https://www.10xgenomics.com/resources/datasets/human-breast-cancer-block-a-section-2-1-standard-1-1-0, June 2020. [Online; accessed 17-April-2023].

[80] 10x Genomics. Human Heart. Available at https://www.10xgenomics.com/resources/datasets/human-heart-1-standard-1-1-0, June 2020. [Online; accessed 17-April-2023].

[81] 10x Genomics. Human Lymph Node. Available at https://www.10xgenomics.com/resources/datasets/human-lymph-node-1-standard-1-1-0, June 2020. [Online; accessed 17-April-2023].

[82] 10x Genomics. Aggregate of mouse brain sections: Visium fresh frozen, whole transcriptome. Available at https://www.10xgenomics.com/resources/datasets/aggregate-of-mouse-brain-sections-visium-fresh-frozen-whole-transcriptome-1-standard, July 2022. [Online; accessed 17-April-2023].

[83] Kristen R Maynard, Leonardo Collado-Torres, Lukas M Weber, Cedric Uytingco, Brianna K Barry, Stephen R Williams, Joseph L Catallini, Matthew N Tran, Zachary Besich, Madhavi Tippani, et al. Transcriptome-scale spatial gene expression in the human dorsolateral prefrontal cortex. Nature neuroscience, 24(3):425–436, 2021.

[84] Yang Liu, Mingyu Yang, Yanxiang Deng, Graham Su, Archibald Enninful, Cindy C Guo, Toma Tebaldi, D. Zhang, Dongjoo Kim, Zhiliang Bai, et al. High-spatial-resolution multi-omics sequencing via deterministic barcoding in tissue. Cell, 183(6):1665–1681, 2020.

[85] Shanshan He, Ruchir Bhatt, Carl Brown, Emily A Brown, Derek L Buhr, Kan Chantranuvatana, Patrick Danaher, Dwayne Dunaway, Ryan G Garrison, Gary Geiss, et al. High-plex imaging of rna and proteins at subcellular resolution in fixed tissue by spatial molecular imaging. Nature Biotechnology, 40(12):1794–1806, 2022.

[86] Chee-Huat Linus Eng, Michael Lawson, Qian Zhu, Ruben Dries, Noushin Koulena, Yodai Takei, Jina Yun, Christopher Cronin, Christoph Karp, Guo-Cheng Yuan, et al. Transcriptome-scale super-resolved imaging in tissues by rna seqfish+. Nature, 568(7751):235–239, 2019.

[87] Patrik L Stahl, Fredrik Salmen, Sanja Vickovic, Anna Lundmark, Jose Fernandez Navarro, Jens Magnusson, Stefania Giacomello, Michaela Asp, Jakub O Westholm, Mikael Huss, et al. Visualization and analysis of gene expression in tissue sections by spatial transcriptomics. Science, 353(6294):78–82, 2016.

[88] Michaela Asp, Stefania Giacomello, Ludvig Larsson, Chenglin Wu, Daniel Furth, Xiaoyan Qian, Eva Wardell, Joaquin Custodio, Johan Reimegard, Fredrik Salmen, et al. A spatiotemporal organ-wide gene expression and cell atlas of the developing human heart. Cell, 179(7):1647–1660, 2019.

[89] Kim Thrane, Hanna Eriksson, Jonas Maaskola, Johan Hansson, and Joakim Lundeberg. Spatially resolved transcriptomics enables dissection of genetic heterogeneity in stage iii cutaneous malignant melanoma. Cancer research, 78(20):5970–5979, 2018.

[90] Jason D Buenrostro, M Ryan Corces, Caleb A Lareau, Beijing Wu, Alicia N Schep, Martin J Aryee, Ravindra Majeti, Howard Y Chang, and William J Greenleaf. Integrated single-cell analysis maps the continuous regulatory landscape of human hematopoietic differentiation. Cell, 173(6):1535–1548, 2018.

[91] 10X Genomics. Fresh cortex, hippocampus, and ventricular zone from embryonic mouse brain (e18). Available at https://www.10xgenomics.com/resources/datasets/fresh-cortex-hippocampus-and-ventricular-zone-from-embryonic-mouse-brain-e-18-1-standard-1-2-0, November 2019. [Online; accessed 17-April-2023].

[92] 10X Genomics. 10k peripheral blood mononuclear cells (pbmcs) from a healthy donor. Available at https://www.10xgenomics.com/resources/datasets/10-k-peripheral-blood-mononuclear-cells-pbm-cs-from-a-healthy-donor-1-standard-1-2-0, November 2019. [Online; accessed 17-April-2023].

[93] 10X Genomics. 8k adult mouse cortex cells, atac v1.1, chromium x. Available at https://www.10xgenomics.com/resources/datasets/8k-adult-mouse-cortex-cells-atac-v1-1-chromium-x-1-1-standard, March 2022. [Online; accessed 17-April-2023].

[94] 10X Genomics. 10k human pbmcs, atac v1.1, chromium x. Available at https://www.10xgenomics.com/resources/datasets/10k-human-pbmcs-atac-v1-1-chromium-x-1-1-standard, March 2022. [Online; accessed 17-April-2023].

[95] Shaun Purcell, Benjamin Neale, Kathe Todd-Brown, Lori Thomas, Manuel AR Ferreira, David Bender, Julian Maller, Pamela Sklar, Paul IW De Bakker, Mark J Daly, et al. Plink: a tool set for whole-genome association and population-based linkage analyses. The American journal of human genetics, 81(3):559–575, 2007.

[96] Frederique Ruf-Zamojski, Zidong Zhang, Michel Zamojski, Gregory R Smith, Natalia Mendelev, Hanqing Liu, German Nudelman, Mika Moriwaki, Hanna Pincas, Rosa Gomez Castanon, et al. Single nucleus multi-omics regulatory landscape of the murine pituitary. Nature communications, 12(1): 2677, 2021.

[97] Hanqing Liu, Jingtian Zhou, Wei Tian, Chongyuan Luo, Anna Bartlett, Andrew Aldridge, Jacinta Lucero, Julia K Osteen, Joseph R Nery, Huaming Chen, et al. Dna methylation atlas of the mouse brain at single-cell resolution. Nature, 598(7879):120–128, 2021.

[98] Haley M Amemiya, Anshul Kundaje, and Alan P Boyle. The encode blacklist: identification of problematic regions of the genome. Scientific reports, 9(1): 9354, 2019.

[99] Simon Peron, Nicholas Sofroniew, Karel Svoboda, Adam Packer, Lloyd Russell, Michael Hausser, Jeff Zaremba, Patrick Kaifosh, Attila Losonczy, Selmaan Chettih, Matthias Minderer, Chris Harvey, Maxwell Rebo, Matthew Conlen, and Jeffrey Freeman. neurofinder: benchmarking challenge for finding neurons in calcium imaging data. Available at https://github.com/codeneuro/neurofinder, March 2016. [Online; accessed 02-January-2024].

[100] Timothy M Johanson, Hannah D Coughlan, Aaron TL Lun, Naiara G Bediaga, Gaetano Naselli, Alexandra L Garnham, Leonard C Harrison, Gordon K Smyth, and Rhys S Allan. Genome-wide analysis reveals no evidence of trans chromosomal regulation of mammalian immune development. PLoS genetics, 14(6):e1007431, 2018.

[101] Naiara G Bediaga, Hannah D Coughlan, Timothy M Johanson, Alexandra L Garnham, Gaetano Naselli, Jan Schroder, Liam G Fearnley, Esther Bandala-Sanchez, Rhys S Allan, Gordon K Smyth, et al. Multi-level remodelling of chromatin underlying activation of human t cells. Scientific reports, 11(1): 528, 2021.

