## Supplementary texts and figures for "Principled PCA separates signal from noise in omics count data"

### S1 Supplementary Methods

#### S1.1 Reparametrization of the Quadratic Variance Function

Non-negativity constraints in $b$ and $c$ in Equation (11) allow further simplification of our objective. Substituting $a = 0$, $b = \sigma^2(1-q)/(1-\sigma^2 q)$, and $c = \sigma^2 q/(1-\sigma^2 q)$ in Equation (7) for any $q_* \in [0,1]$ and $\sigma > 0$, gives

$$\widehat{\mathrm{Var}}[Y_{ij}] = \sigma^2 \left[(1-q)Y_{ij} + qY_{ij}^2\right] . \tag{19}$$

This formulation imposes that the data variance is a convex combination of linear and quadratic variance scaled by the noise variance $\sigma^2$. As noted in [36], because $\sigma$ is a global scaling factor, the biwhitening factors of a given data matrix $Y$ are determined by the choice of $q_* \in [0,1]$. That is, if $(\hat{\boldsymbol{u}}_{*,1}, \hat{\boldsymbol{v}}_{*,1})$ are the biwhitening factors learned by substituting $q_*$ and $\sigma = 1$ into Equation (19), then $(\sigma_*^{-1/2}\hat{\boldsymbol{u}}_{*,1}, \sigma_*^{-1/2}\hat{\boldsymbol{v}}_{*,1})$ are the biwhitening factors for any $\sigma_* > 0$. Consequently, the biwhitened data matrix for any pair $q_*$ and $\sigma_*$, $\widetilde{Y}_{q_*,\sigma_*}$, is $\sigma_*^{-1}\widetilde{Y}_{q_*,1}$.

For a given $q_*$, an optimal $\sigma_*$ can be estimated by matching a quantile $p \in [0,1]$ of the empirical spectral distribution of $\hat{\sigma}_*^{-2}\widetilde{\Sigma}_{q_*,1}$ to the corresponding quantile of a unit MP distribution. That is,

$$\hat{\sigma}(q_*) = \hat{\sigma}_* = \frac{s_{\geq \lceil pm \rceil}(\widetilde{Y}_{q_*,1})}{\sqrt{n\mu_{p,\gamma}}} , \tag{20}$$

where $\gamma = m/n$, $s_{\geq \lceil pm \rceil}(\widetilde{Y}_{q_*,1})$ is the smallest singular value of $\widetilde{Y}_{q_*,1}$ that is greater than or equal to $\lceil pm \rceil$ singular values of $\widetilde{Y}_{q_*,1}$, and $\mu_{p,\gamma}$ is the $p$-th quantile of the MP distribution with aspect ratio $\gamma$ and noise variance 1, i.e., $\mu_{p,\gamma}$ is the solution to the equation

$$F_{\gamma,1}(t) = \int_{\beta_-}^{t} \frac{\sqrt{(\tau - \beta_-)(\beta_+ - \tau)}}{2\pi\gamma\tau} d\tau = p,$$

in the variable $t \in (\beta_-, \beta_+)$, and $\beta_\pm = (1 \pm \sqrt{\gamma})^2$. Our definition is a small modification of the noise level estimator described by [40, 39]; it is equivalent when $p = 0.5$. In that case, which we first described in [36], the median of the biwhitened data spectrum is matched to the median of the MP. There, we noted that the median is an advantageous choice for quantile scaling because it is robust to outlier eigenvalues (e.g., those attributable to signal). In this work, we allow $p$ to be any quantile, which trades robustness in $\sigma$ estimation for computational considerations, as computing $\hat{\sigma}(q_*)$ for $p > 0.5$ requires computing fewer singular values of $\widetilde{Y}_{q_*,1}$.

In summary, we learn QVF parameters for a dataset by minimizing the discrepancy between the empirical spectral distribution of the biwhitened data and the MP distribution. In this work, we limit our search to distributions that have zero constant variance and non-negative linear and quadratic variance. That is, we assume $a = 0$ and learn estimates $\hat{b} \geq 0$ $\hat{c} \geq 0$ by solving problem Equation (11). Subsequently, we re-parameterize $b$ and $c$ such that they satisfy a convex combination with coefficient $q$ up to global scaling by $\sigma^2$ Equation (19). This is an optimization in a single variable, $q$, as our estimate of $\sigma$ is a function of $q$ derived by estimating the noise level of the data given $q$. That is, we estimate $q$ by solving

$$\begin{aligned}
\hat{q} = \arg\min_{q_*} \quad & \mathrm{KS}_{\mathrm{MP},Y}(0, b_*, c_*) \\
\text{s.t.} \quad & q_* \in [0,1] \\
& b_* = \hat{\sigma}(q_*)^2(1-q_*)/(1-\hat{\sigma}(q_*)^2 q_*) \\
& c_* = \hat{\sigma}(q_*)^2 q_*/(1-\hat{\sigma}(q_*)^2 q_*) ,
\end{aligned} \tag{21}$$

and estimate $\sigma$ by $\hat{\sigma}(\hat{q})$.

#### S1.2 Optimization implementation

Our implementation allows flexibility in the choice of one's solver. In this work, and by default in our implementation, we use polynomial approximation using Chebyshev polynomials [64]. Polynomial approximation features strong stability guarantees and easily computable bounds on approximation errors. Furthermore, for a given matrix and approximation degree, polynomial approximations proceed in constant time, and one can vary the approximation degree to trade between accuracy and speed. In practice, we find that the approximation error of Equation (21) using $d = 64$ is typically at most $10^{-6}$, which leads to visibly good fits of biwhitened data to the MP.

Given a solution $\hat{q}$ for Equation (21), we obtain estimates for $\hat{b}$ and $\hat{c}$ by computing $\hat{\sigma}(\hat{q})$. This can be computed directly given the singular values of $\tilde{Y}_{\hat{q},1}$ or through polynomial approximation. Because every evaluation in solving Equation (21) also requires evaluating Equation (20), we can approximate Equation (20) without any extra computations.

#### S1.3 Computational considerations: efficiency

BiPCA employs two costly algorithms: Sinkhorn-Knopp matrix scaling and singular value decomposition (SVD). For this work, we have chosen to direct our efforts towards reducing the number and size of SVDs used by BiPCA. We did this because, in practice, Sinkhorn-Knopp matrix scaling is not the bottleneck of BiPCA.

Sinkhorn-Knopp matrix scaling requires $O(tmn)$ computations to proceed for $t$ iterations on an $m \times n$ matrix. In practice, we find that Sinkhorn-Knopp matrix scaling typically converges quickly: by default, our implementation proceeds until a maximum scaling error of $10^{-6}$ is met, which often occurs in less than 100 iterations. On the other hand, a complete SVD of an $m \times n$ matrix $Y$ requires $O(m^2 n)$ operations. Furthermore, if the data matrix $Y$ is large and sparse, complete SVD will be infeasible due to memory constraints. Finally, we note that our pipeline uses a degree $d$ Chebyshev polynomial approximation to learn QVF coefficients - naively, this requires $d+1$ complete SVDs of an $m \times n$ matrix. There are a few ways to address the SVD requirements of BiPCA.

**Low-rank assumptions allow truncated singular value decompositions.** Since we assume the signal contained in $\widetilde{Y}$ is low-rank and contained in $r$ singular vectors with scaled singular values greater than the MP upper edge $\beta_+$, we only need to compute a small subset of singular values to denoise the data. There are two cases where this is applicable: 1) if we have a reliable estimate of its biwhitening factors; or 2) if $\hat{q}$ is known but $\hat{\sigma}$ is not, and we tolerate instabilities in its estimation. In these cases we need only compute $r$ singular values of $\widetilde{Y}$ to perform denoising. In the second case, we can compute $\lceil(1-p)m + r\rceil$ singular values of $\widetilde{Y}$. Truncated and randomized SVD algorithms can be used in this situation, and they are effective for sparse matrices. In practice, this involves computing slightly more than $r$ ($\lceil(1-p)m+r\rceil$, respectively) singular values. Since $r$ is unknown, we must compute more than $r$ ($\lceil(1-p)m+r\rceil$, resp.) singular values to ensure that a) $\beta_+$ is found and b) the first $r$ ($\lceil(1-p)m+r\rceil$, resp.) signal singular values are computed accurately (the accuracy of truncated and particularly randomized SVD algorithms can be improved by over-estimating the rank of the matrix). In this situation, by default our pipeline iteratively computes singular values until $\beta_+$ is surpassed (or the $p$-th quantile of the singular values is obtained). We recommend this approach when QVF parameters are known and the data matrix is sparse, such as in SNP data.

**QVF parameters can be estimated for large matrices using random sampling.** If QVF parameters are unknown, they must be estimated. If the data matrix is large, then the optimization described in previous sections will be infeasible. For this situation we implemented a constant-time solution to estimate QVF parameters. Our approach is based on the observation that the QVF coefficients $a$, $b$, and $c$ do not vary over the elements of the data matrix. Because the same QVF parameters are used for every submatrix of the data, we assume that we can approximate our estimates by choosing $\hat{b}$ and $\hat{c}$ that minimize $\mathrm{KS}_{\mathrm{MP},Y_{IJ}}(0, \hat{b}, \hat{c})$ *on average*, where $Y_{IJ}$ is a randomly drawn submatrix of $Y$ with row index set $I \subset \{1, \ldots, m\}$ and column index set $J \subset \{1, \ldots, n\}$. Letting $m_s$ be a user-designated size of $I$ and $J$, $n_s$ be a user-designated number of submatrices sampled, and $d$ be a user-designated approximation degree, then this approach requires $O(n_s m_s^3 [d+1])$ operations. As none of these parameters scale with the size of the input data, QVF coefficient fitting in this way runs in constant time. By default, we choose $m_s = \min\{5000, m\}$, $n_s = 5$, and $d = 64$. In practice, we found that these choices represent a reasonable trade off between variance in approximating $\hat{b}$ and $\hat{c}$ at the expense of computation time.

**Random sampling and truncated singular value decomposition can be combined.** For settings where a matrix is large but can be stored in memory, i.e., those in which computing one complete SVD is feasible but $d+1$ SVDs is intractible, we recommend using random sampling to estimate QVF parameters before subsequently applying a complete SVD. This ensures the accuracy of singular vectors, singular values, and estimation of noise variance by $\hat{\sigma}$.

For settings where computing a single complete SVD is intractable, then random sampling can be performed to estimate QVF parameters first before performing truncated SVD. An initial best guess at $\hat{b}$ and $\hat{c}$ can be obtained by polynomial approximation of $\hat{\sigma}$. Subsequently, one can iteratively compute singular values (as previously described) until $\beta_+$ is met. If enough singular values are obtained to perform noise variance estimation using the $p$-th quantile, $\hat{b}$ and $\hat{c}$ can be further refined by computing $\hat{\sigma}$. We emphasize that this refinement will be sensitive to the choice of $p$ and the rank $r$ - whereas the median is robust in the low rank matrices we consider in this paper, extreme quantiles of the empirical spectrum (such as those that would be feasible to compute using randomized SVD) will be sensitive. In practice, we find that approximation of $\hat{\sigma}$ typically performs well enough that any final steps of $\hat{\sigma}$ refinement provide marginal benefits.

### S2 Details of Data Processing

Here we provide the details regarding processing each dataset used in the study. The size of each dataset before and after filtering is detailed in Supplementary table 1.

#### S2.1 Single-cell or single-nucleus RNA sequencing

**Smart-seq3** *HagemannJensen2020*: We analyzed the Human Cell Atlas (HCA) dataset from [67]. HEK cells are removed from the data. We removed cells that have less than 75% mapped reads, total read number $\leq 10^5$, and cells that have less than 500 genes expressed following [67]. We further removed genes that have less than 250 cells expressed to ensure the matrix is not too sparse to run BiPCA.

**Smart-seq3xpress** *HagemannJensen2022*: We analyzed the Human PBMC dataset from [68]. We first split the data by donors (7 donors). Then following [68], we removed cells that (1) have less than 50% mapped reads, or (2) have total number of reads $< 20,000$, or (3) more than 15% reads mapped to mitochondrial genes. In addition, we imposed a sparsity filter to remove cells that have less than 500 genes expressed and genes that have less than 50 cells expressed.

**10X Genomics Chromium V1** *10X2016PBMC*: We analyzed the human PBMC dataset from [66]. We obtained the filtered count matrix from 10x Genomics website. We further removed cells that have less than 100 genes expressed and cells that have more than 10% UMIs mapped to mitochondrial genes. We also removed genes that have less than 100 cells expressed.

*Zheng2017*: We analyzed the human PBMC dataset from [49]. Filtered expression matrices were obtained for FACS purified CD19+ B cells, CD4+ helper T cells, CD14+ cells, CD34+ cells, CD4+CD25+ regulatory T cells, CD4+CD45RA+CD25- naive T cells, CD4+CD45RO+ memory T cells, CD56+ natural killer (NK) cells, CD8+ cytotoxic T cells, and CD8+CD45RA+ naive cytotoxic T cells from the 10X Genomics website. CD14+ cells were further split into CD14+ monocytes and Dendritic cells (DCs). The matrices were then merged. Further processing steps for the analysis (marker gene analysis and rank estimation experiment) are detailed in each section.

We also analyzed each cell type separately as individual datasets where we run BiPCA on each count matrix. For each matrix, we filter out cells that have less than 100 genes expressed and more than 10% UMIs mapped to mitochondrial genes. We also filter out genes that have less than 100 cells expressed.

**10X Genomics Chromium V2** *10X2017MouseBrain*, *10X2017PBMC*, *TenX2021HekMixtureV2*: We analyzed the mouse brain dataset from [69], the human PBMC dataset from [70], and the dataset with mixture of Human HEK293T and Mouse NIH3T3 cells from [71]. Each filtered count matrix is obtained from 10x Genomics website. We further removed cells that have less than 100 genes expressed and cells that have more than 10% UMIs mapped to mitochondrial genes. We also removed genes that have less than 100 cells expressed.

**10X Genomics Chromium V3** *10X2018MouseBrain*, *10X2018PBMC*: We analyzed the mouse brain dataset from [72] and the human PBMC dataset from [73]. Each filtered count matrix is obtained from 10x Genomics website. We further removed cells that have less than 100 genes expressed and cells that have more than 10% UMIs mapped to mitochondrial genes. We also removed genes that have less than 100 cells expressed.

*SCORCH_INS*: We analyzed a 10x Chromium single-nucleus RNA-seq dataset (snRNA) of human insular cortex (INS) from the SCORCH consortium [51]. Raw data is first processed through cellranger v6.0.1 using default parameters. The RNA count matrix is then filtered for barcodes with less than 500 non zero entries or more than 7500 non zero entries. Barcodes with more than 2% UMIs mapped to mitochondrial genes are filtered out. We also filter out genes with less than 100 cells expressed. Doublet removal is performed using `scDblFinder` [74] to remove doublets.

**10X Genomics Chromium V3.1** *10X2020MouseBrain*, *TenX2021HekMixtureV3*: We analyzed the mouse brain dataset from [75] and the dataset with mixture of Human HEK293T and Mouse NIH3T3 cells from [76]. We obtained the filtered count matrix from 10x Genomics website. We further removed cells that have less than 100 genes expressed and cells that have more than 10% UMIs mapped to mitochondrial genes. We also removed genes that have less than 100 cells expressed.

*10X2021PBMC*: We analyzed the human PBMC dataset from [77]. We obtained the filtered count matrix from 10x Genomics website. We further removed cells that have less than 100 genes expressed and cells that have more than 10% mitochondrial ratio. We also removed genes that have less than 60 cells expressed.

**CITE-seq (RNA)** *Stoeckius2017*: The RNA count matrix is obtained from [53]. The cell type annotations from the RNA modality and the protein modality are obtained from [78]. We removed mouse genes except the top 100 most highly expressed ones. We removed erythroid cells, megakaryocytes, and dendritic cells as they are not well distinguished in the protein modality. We further removed cells that are not labeled or labeled as doublets from the data. At the end, sparsity filters are applied to remove genes that are detected in fewer than 100 cells and cells that express genes fewer than 100.

*Stuart2019*: The RNA count matrix is obtained from [54]. The cell type annotations are obtained from [78] in which both the RNA and the protein modality are used for annotation [54]. We removed cells that are not annotated. Sparsity filters are applied to remove genes that are detected in fewer than 100 cells and cells that express genes fewer than 100.

*Open-challenge CITE-seq* (*Luecken2021CITE*): The RNA count matrix and the cell type annotations are obtained from [55] across 12 batches. Sparsity filters are applied to each batch to remove genes that are detected in fewer than 100 cells and cells that express genes fewer than 100.

**Multiome (RNA)** *Open-challenge Multiome* (*Luecken2021Multiome*): The RNA count matrix and the cell type annotations are obtained from [55] across 13 batches. Sparsity filters are applied to each batch to remove genes that are detected in fewer than 100 cells and cells that express genes fewer than 100.

*SCORCH_PFC*: Three samples from SCORCH consortium [51], where each sample is collected from human prefrontal cortex (PFC) region and sequenced using 10x Multiome. Raw data is first processed through cellranger-arc v2.0.2 with default parameters. The RNA count matrix is then filtered for barcodes with less than 500 non zero entries or more than 7500 non zero entries. Barcodes with more than 2% mitochondrial genes are filtered out. We also filter out genes with less than 100 cells expressed. Doublet removal is performed using `scDblFinder` [74] to remove doublets.

#### S2.2 Spatial Transcriptomics

**10X Visium v1** Our data collection contained 16 samples obtained from 5 sources.

*10X2020HumanBreastCancer*: Filtered Visium gene expression count matrices samples from two sections of Invasive Ductal Carcinoma breast tissue were obtained from 10X Genomics [79]. Section 1 contained 3,798 spots, and section 2 contained 3,987 spots. The section matrices were concatenated, producing a $7,785$ spots $\times 36,601$ features matrix. $7,785$ spots and $16,276$ features remained after removing spots that have fewer than 20 genes expressed and sparse features detected in fewer than 100 spots.

*10X2020HumanHeart*: A $4,247$ barcode $\times 36,601$ features filtered Visium count matrix sampled from human heart tissue was obtained from 10X Genomics [80]. $4,247$ barcodes and $10,193$ features remained after removing spots that have fewer than 20 genes expressed and sparse features detected in fewer than 100 barcodes.

*10X2020HumanLymphNode*: A $4,035$ barcode $\times 36,601$ features filtered Visium count matrix sampled from a human lymph node was obtained from 10X Genomics [81]. $4,035$ barcodes and $14,901$ features remained after removing spots that have fewer than 20 genes expressed and sparse features detected in fewer than 100 barcodes.

*10X2022MouseBrain*: A $6,112$ barcode $\times 32,285$ features aggregated and filtered Visium count matrix sampled from two sections of mouse brain tissue (Sagittal-Anterior and Sagittal-Posterior) was obtained from 10X Genomics [82]. $6,112$ barcodes and $14,215$ features remained after removing spots that have fewer than 20 genes expressed and sparse features detected in fewer than 100 barcodes.

*Maynard2021*: 12 Visium count matrices of human dorsolateral prefrontal cortex were obtained from [83]. Each unfiltered matrix contained an average of $3,973$ barcodes and $33,538$ features. After sparse feature filtering (features must be detected in $\geq 100$ spots and spots that have $\geq 20$ genes expressed), an average of $10,476$ features remained in each sample.

**Deterministic Barcoding in Tissue for spatial omics sequencing (DBiT-Seq) [84]** We analyzed 1 dataset generated using DBit-Seq.

*Liu2020*: The count matrix is obtained from [84]. It has 2,500 barcodes and 22,969 features. 2,500 barcodes and 10,093 features remained after removing spots that have fewer than 20 genes and sparse features detected in fewer than 100 barcodes.

**nanoString CosMx Spatial Molecular Imager (SMI) [85]** *Kluger2023Melanoma*: We analyzed 1 SMI dataset from a total of 3 different FOVs. To perform BiPCA, each FOV was filtered and analyzed separately. We filter out cells that have less than 100 genes expressed and genes that have less than 100 cell expressed.

*FrontalCortex6k*: We analyzed 1 SMI dataset from human frontal cortex [48]. This dataset includes in total 194,065 cells and 6,078 gene targets, spanning over 392 FOVs. We focused on FOV 63 that has the most cells (960), and we further filter out cells that have less than 100 genes expressed and genes that have less than 1 cell expressed. In total, we have 907 cells and 6,256 genes left after filtering.

**Sequential fluorescence in situ hybridization (seqFISH+)** [86]

*Eng2019*: We analyzed the seqFISH+ data of 2 regions (sub-ventricular zone (SVZ) and olfactory bulb) from [86], where SVZ has 913 cells and 10,000 features and olfactory bulb has 2,050 cells and 10,000 features. After filtering out features that are detected in fewer than 100 cells and cells that have fewer than 20 features expressed, the number of remaining features is 9,566 for SVZ and 5,769 for olfactory bulb.

**Spatial Transcriptomics V1.0.0 [87]** *Asp2019*: The count matrix is obtained from [88]. It has 3,111 barcodes and 39,739 features. 3,111 barcodes and 11,260 features remained after removing barcodes that have fewer than 20 features expressed and sparse features detected in fewer than 100 barcodes.

*Thrane2018*: The dataset is obtained from [89] . We analyzed 8 datasets (4 samples of stage III melanoma lymph node metastases, 2 sections each) from [89]. We removed barcodes that have fewer than 20 features expressed and sparse features detected in fewer than 100 barcodes.

#### S2.3 Single-cell or single-nucleus ATAC sequencing

**Fluidigm C1 [43]** *Buenrostro2018*: We analyzed the human hematopoietic cells dataset from [90]. The fragment count data is filtered to exclude cells with less than 1000 non-zero features (peaks) and features (peaks) that have non-zero values in less than 100 cells.

**10X Genomics Single Cell ATAC V1** *10X2019MouseBrainATAC* and *10X2019PBMCATAC*: We analyzed the mouse brain dataset from [91] and the human PBMC dataset from [92]. Each filtered peak by cell count matrix is obtained from 10x Genomics website and is filtered to exclude cells with less than 1000 non-zero features (peaks) and features (peaks) that have non-zero values in less than 100 cells.

**10X Genomics Single Cell ATAC V1.1** *10X2022MouseCortexATAC* and *10X2022PBMCATAC*: We analyzed the mouse cortex dataset from [93] and the human PBMC dataset from [94]. Each filtered peak by cell count matrix is obtained from 10x Genomics website and is filtered to exclude cells with less than 1000 non-zero features (peaks) and features (peaks) that have non-zero values in less than 100 cells.

**Multiome (ATAC)** *Open-challenge Multiome* (*Luecken2021Multiome*): The ATAC count matrix and the cell type annotations are obtained from [55] across 13 batches. Sparsity filters are applied to each batch to remove peaks that have non-zero values in less than 150 cells and cells with less than 100 non-zero peaks. For the experiment on preserving cell neighborhoods, we remove peaks that are detected in less than 50 cells in order to keep more informative peaks in the experiment.

#### S2.4 Other count modalities

**1000 Genome Genotyping Data** *Byrska2022*: We analyzed the phased 1000 Genome phase3 data [3] where 3,202 samples were genotyped in total. We use Plink [95] to preprocess the data as follows: We only included chromosomes 1-22 in the analysis and we filtered out variants with more than 2 alleles and variants with minor allele frequency below 0.1. Additionally, we removed 630 samples with first and second degree relations, and pruned the variants that are in approximate linkage equilibrium with each other with the following parameters: window size =

200, shift = 5, and variance inflation factor = 1.005. After the Plink preprocessing steps, we obtained a genotype data matrix with $2,573$ samples and $17,840$ variants.

**Single Cell Methylomics Data** *RufZamojski2021*: We analyzed snmC-seq2 data of the adult (age P56) male mouse brain pituitary (PIT) from [96], following the pipeline provided by [97]. We started with the ALL Cytosine (ALLC) file, which contains base-level methylation and coverage counts for each cell. This file was processed and aggregated by ALLCools v1.0.21[97] into an MCDS file containing two count matrices for cells $\times$ 5kb genomic bins: one for methylated CG counts and another for total CG counts. We processed these two count matrices together. We excluded cells using the suggested cutoffs provided by [97], specifically cells with a mapping rate of less than 0.5, fewer than $500,000$ final mC reads, an mCCC fraction larger than 0.03, an mCH fraction larger than 0.2, and an mCG fraction of less than 0.5. Next, we filtered out bins overlapping $> 0.2$ with ENCODE mm10 blacklist regions v2[98] and bins in chromosomes M and Y. To maintain the aspect ratio of rows and columns at less than 5, we randomly sampled the bins to five times the number of cells. We stabilized the matrix by filtering out bins with fewer than 5 cells expressed and cells with fewer than 5 bins expressed. In total, we obtained a cells $\times$ bins matrix of $2,756 \times 13,656$ for both the methylated CG count matrix and the total CG count matrix.

**Calcium Imaging Data** *Neurofinder2016*: We analyzed one sample from [99] with $8,000$ calcium images of neurons. Each image is $512 \times 512$. We flattened the images and concatenate all images together for running BiPCA.

**Hi-C Data** *Johanson2018*: We analyzed 10 samples from two data sources [100, 101] that consist of native CD4 T cells, naive CD8 T cells, B cells, activated CD4 T cells, and activated CD8 T cells from two human subjects. We select only on the interactions between chromosome 1 and chromosome 2 and filter out regions on each chromosome that have less than 10 total contacts with the other chromosome.

### S3 Supplementary Figures

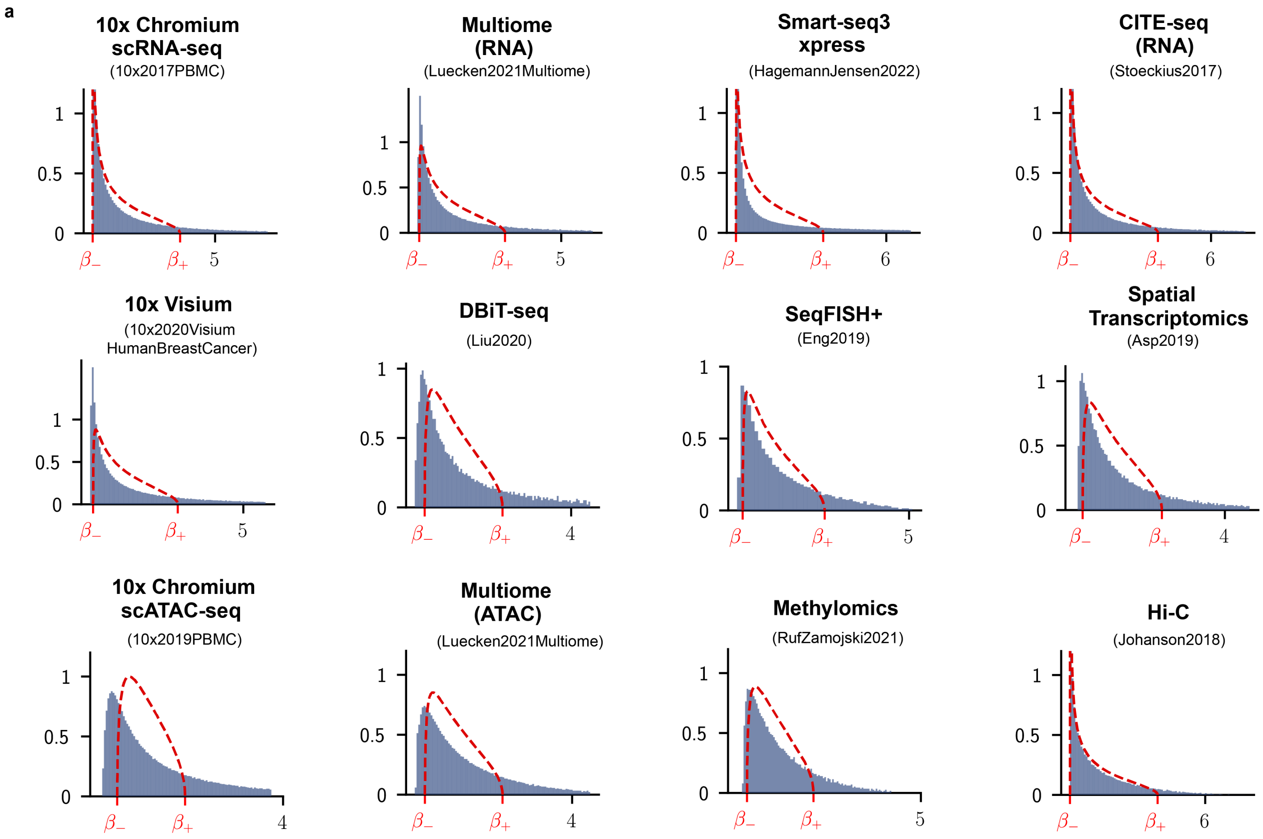


**Figure S1:** Unnormalized data do not fit the MP distributions. Each panel shows the scaled empirical spectral distribution of an example dataset from each modality (blue histogram), whereas the red dashed curve is the theoretical MP distribution. The empirical spectral distributions are scaled by a constant for visualization.

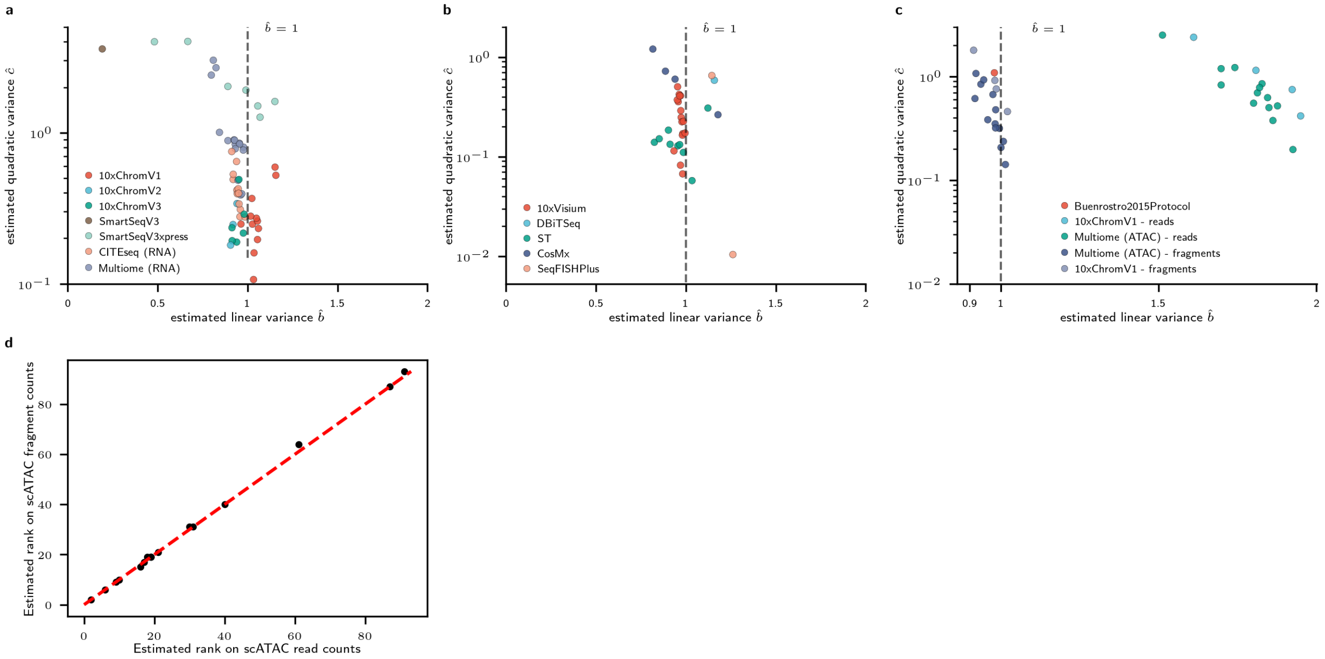


**Figure S2:** Analysis of BiPCA parameter estimates. **a-c**: Visualization of the estimated b (x-axis) and c (y-axis) parameters across datasets from scRNA-seq (**a**), spatial transcriptomics (**b**), and scATAC-seq (**c**). Each dataset is colored by the corresponding protocol. **d**: Estimated rank for the ATAC read counts (x-axis) and the ATAC fragment counts (y-axis) for each scATAC-seq dataset in the compendium (black points). A point that lies on the red dashed line represents the estimated rank is the same between two matrices.

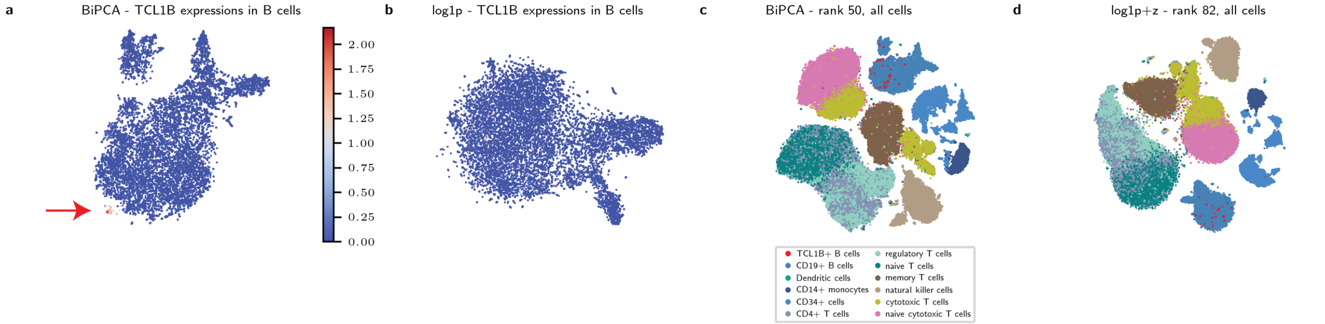


**Figure S3:** T-SNE embeddings of the *zheng2017* PBMC dataset. (**a**) Visualization of the TCL1B expression levels on the t-SNE embeddings of B cells from BiPCA. Red arrow highlights the TCL1B+ B cell population. (**b**) Visualization of the TCL1B expression levels on the t-SNE embeddings of B cells from log1p+z. (**c**) Visualization of different cell types on the t-SNE embeddings with 50 PCs from BiPCA as input. (**d**) Visualization of different cell types on the t-SNE embeddings with 82 PCs from log1p+z as input.

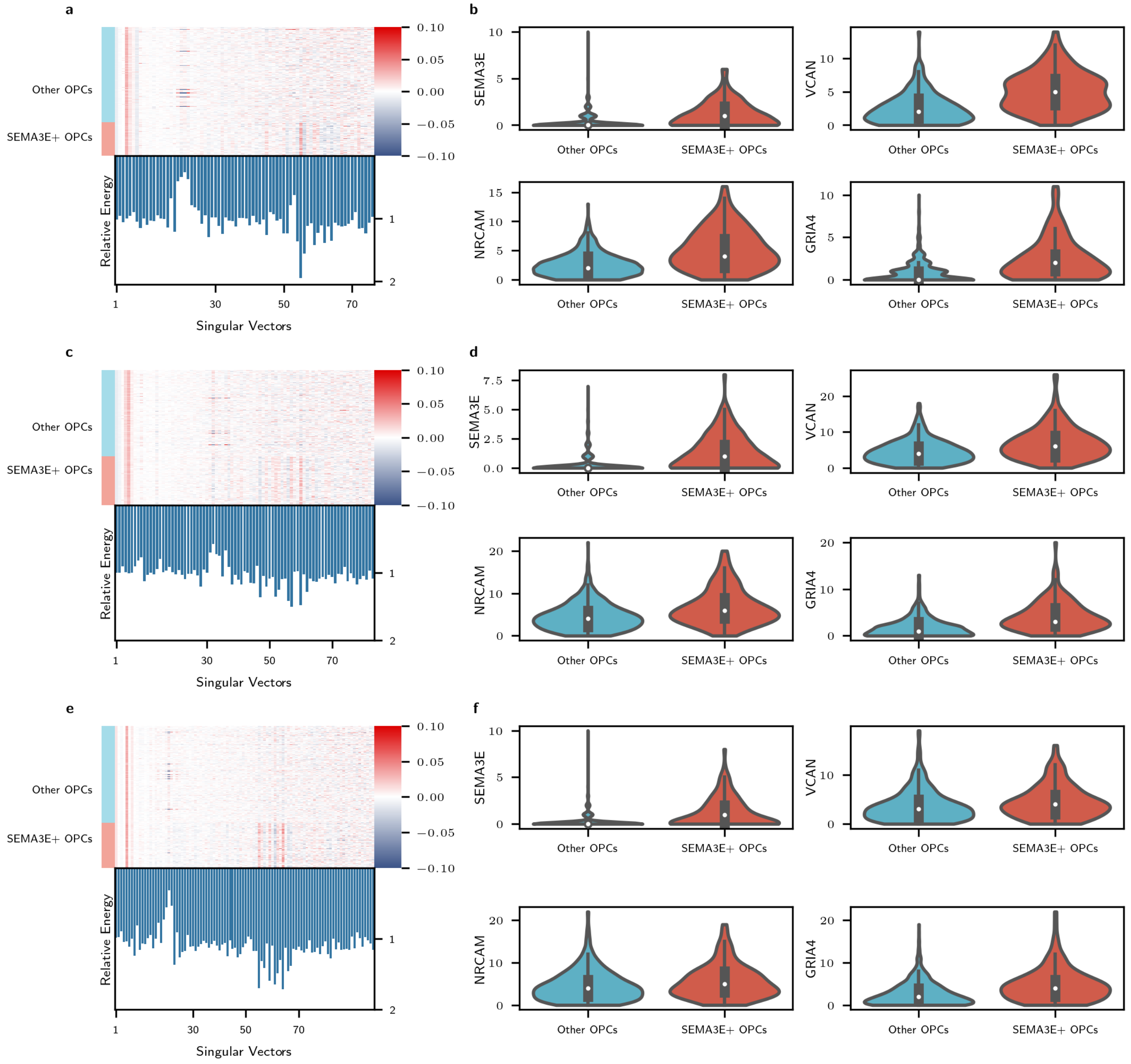


**Figure S4:** Identification of the SEMA3E+ oligodendrocyte precursor cells (OPCs) in three 10x Multiome human prefrontal cortex (PFC) samples. (**a-b**,**c-d**,**e-f** for each sample separately).**a** Top: heatmap visualization of the signal components (singular vectors) across OPCs for the human PFC data. The columns are singular vectors ordered by singular values, and the rows are OPCs groups by SEMA3E+ OPCs and Other OPCs. Bottom: Relative energy of the identified OPC cluster within all OPCs for each singular vector. The singular vectors with the largest relative energy are indexed at: 55 and 56. **b**: Violin plot visualization of the top marker genes for the identified OPC cluster. **c-f**: Validation of the identified SEMA3E+ OPCs in two additional samples (**c-d** for the 2nd sample, **e-f** for the 3rd sample).

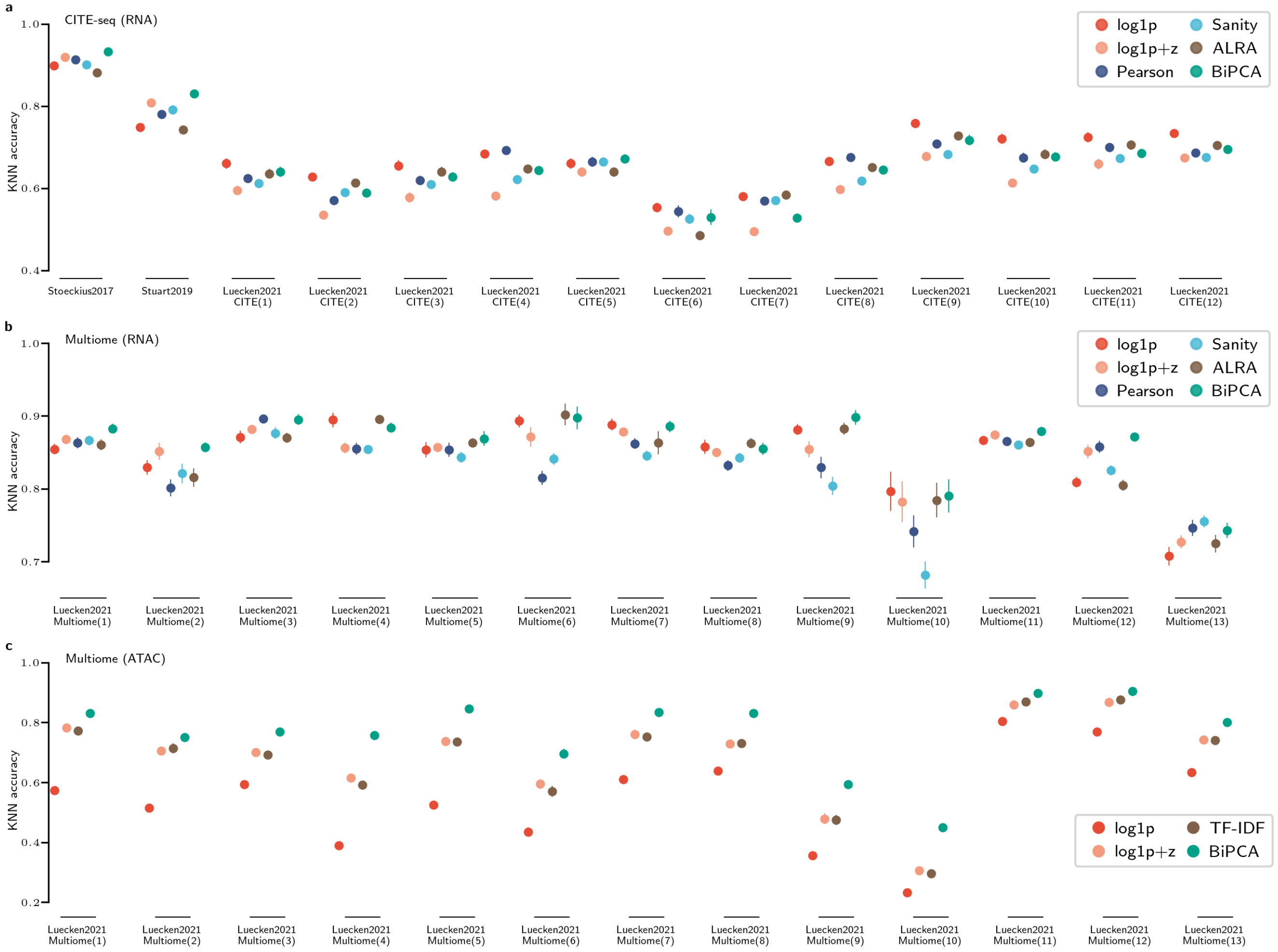


**Figure S5:** Post-normalization k-NN cell type classification accuracy in 3 modalities. (**a**) CITE-seq (RNA): the RNA counts of 14 CITE-seq datasets were used to classify cell types assigned by surface protein labels; (**b**) Multiome (RNA): RNA counts from 13 Multiome datasets were used to classify cell types annotated using ATAC counts; (**c**) Multiome (ATAC): ATAC counts from the 13 Multiome datasets in **b** were used to classify cell types annotated using RNA counts. Each experiment is repeated 10 times with 80% of the data. Within each dataset, different normalization methods are applied (colored dots) and accuracies are computed on the PCA space in each method. Mean accuracy and the standard deviations are plotted.

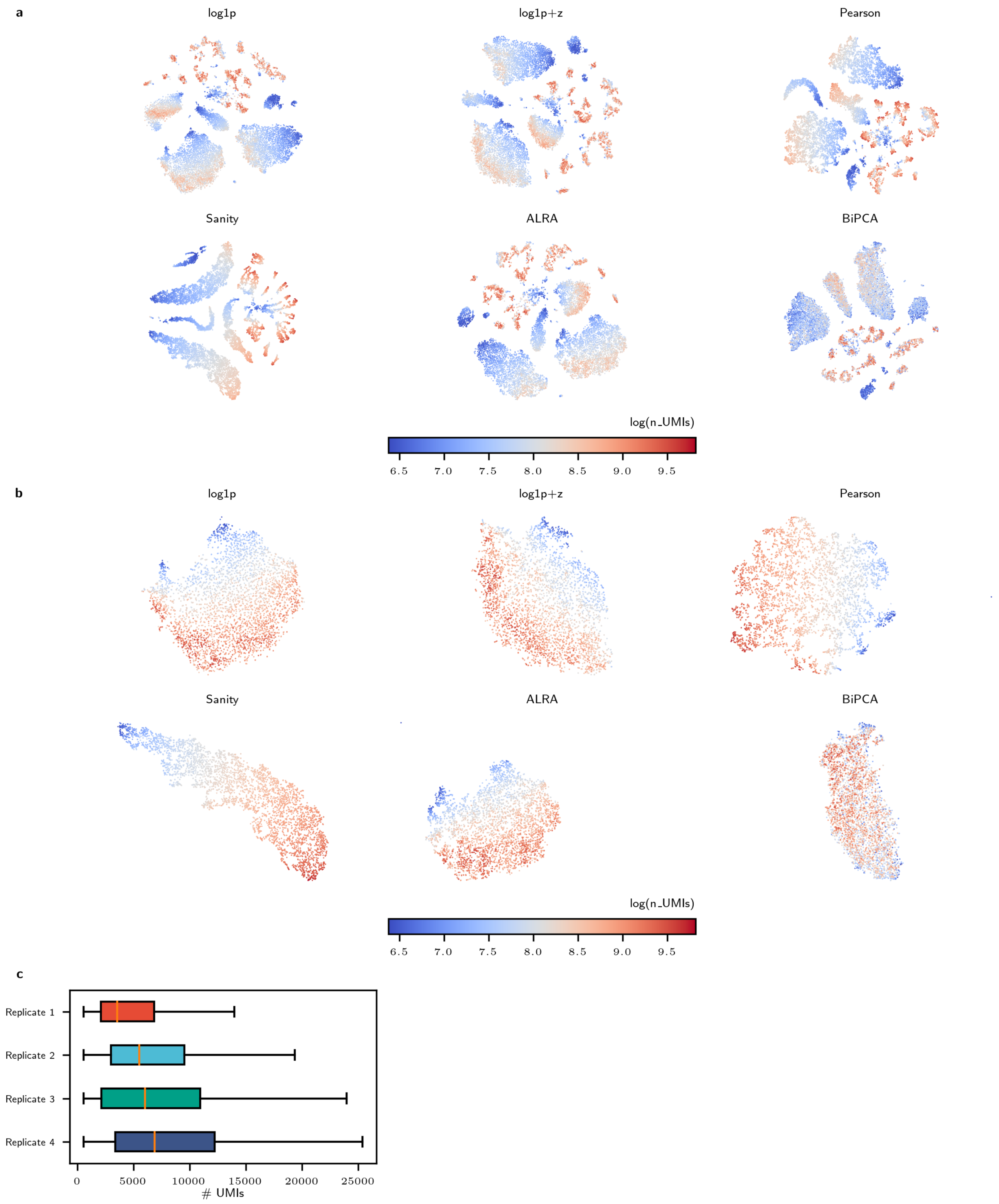


**Figure S6:** Library size effects in *Kluger2024OUD*. A: t-SNE visualization after different normalization methods. Cells are colored by the number of UMIs at log scale. B: t-SNE visualization of the astrocytes. C: Distribution of the number of UMIs across replicates.

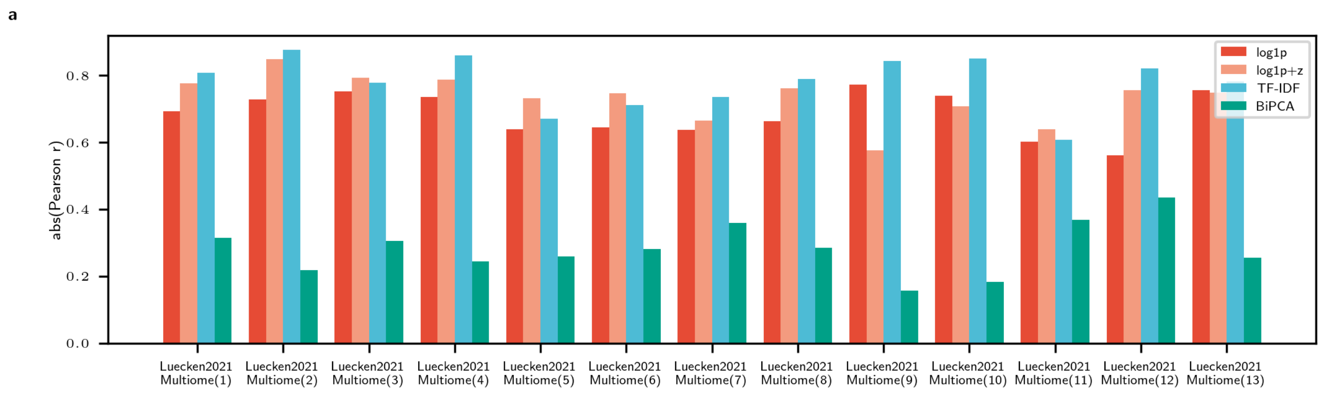


**Figure S7:** Maximum correlation (y-axis) between post-normalization singular vectors and library depth for different methods on the different batches in the *Luecken2021Multiome* [55] (ATAC) data (x-axis).

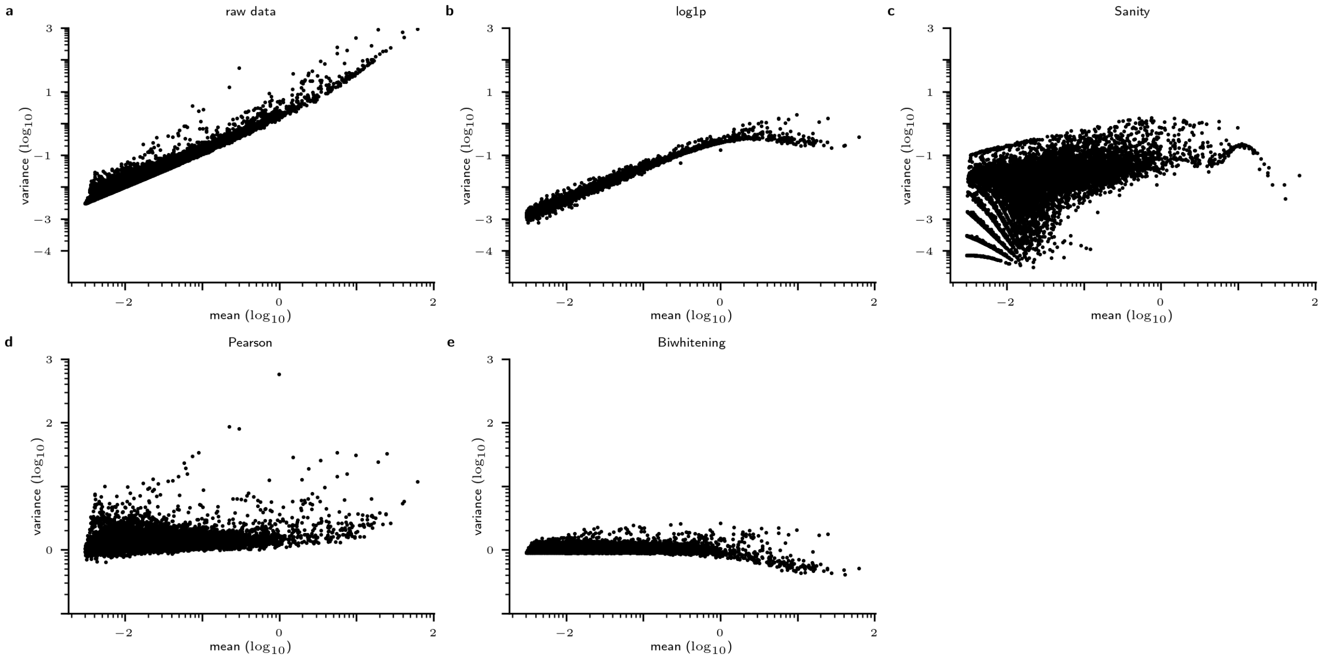


**Figure S8:** Transformed gene variances after methods that do not perform low-rank approximation. Each panel shows the relationship between transformed gene variances (y-axis) relative to its mean expression levels on the raw data (x-axis) for each gene across different transformations: (**a**) raw data, (**b**) log1p, (**c**) Sanity, (**d**) Pearson, (**e**) Biwhitening

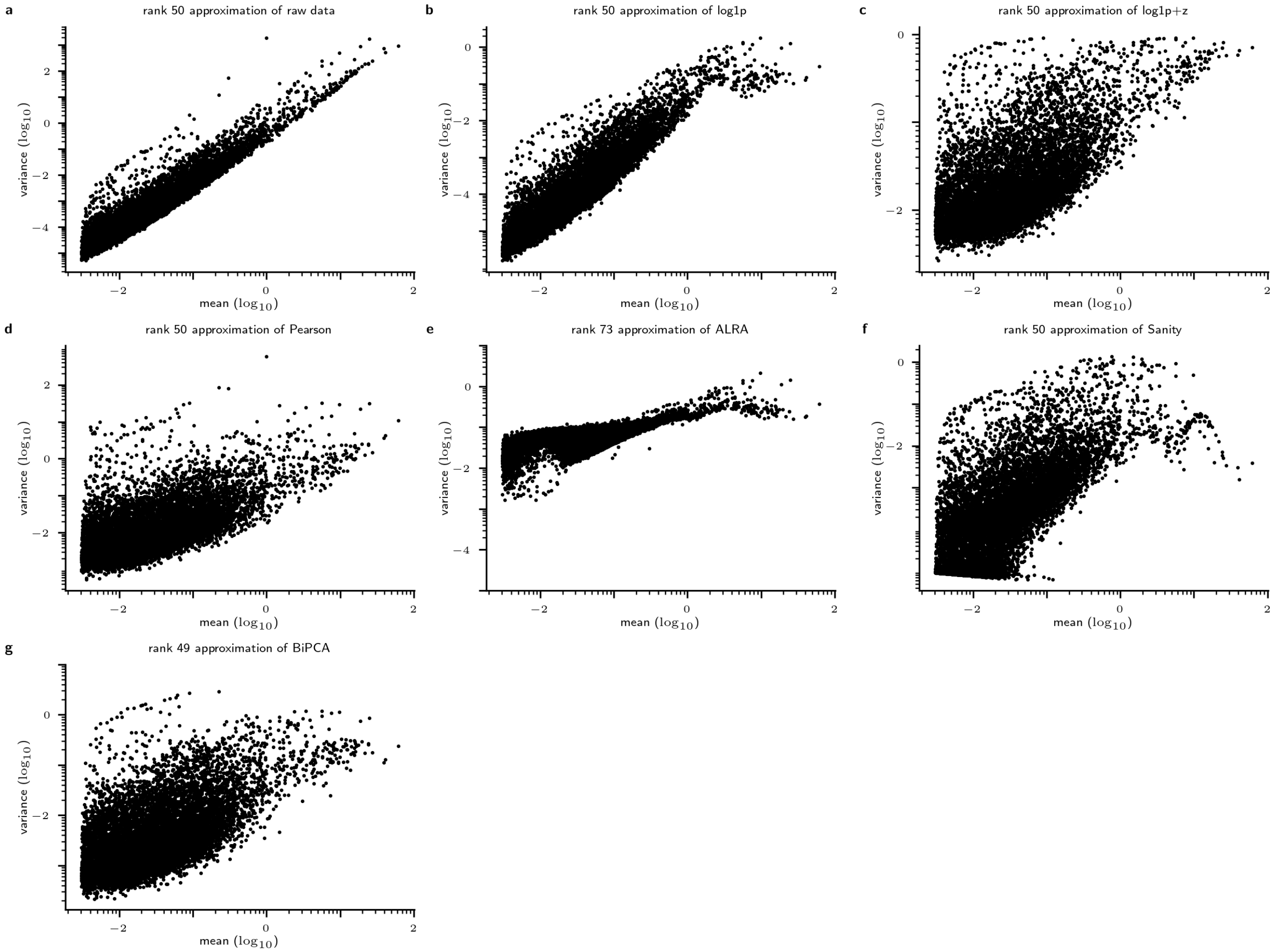


**Figure S9:** Transformed gene variances after methods that perform low-rank approximation. Each panel shows the relationship between transformed gene variances (y-axis) relative to its mean expression levels on the raw data (x-axis) for each gene across different transformations: (**a**) rank 50 approximation of raw data, (**b**) rank 50 approximation of log1p, (**c**) rank 50 approximation of log1p+z, (**d**) rank 50 approximation of Pearson, (**e**) rank 50 approximation of Sanity, (**f**) rank 73 approximation of ALRA, (**g**) rank 49 approximation of BiPCA

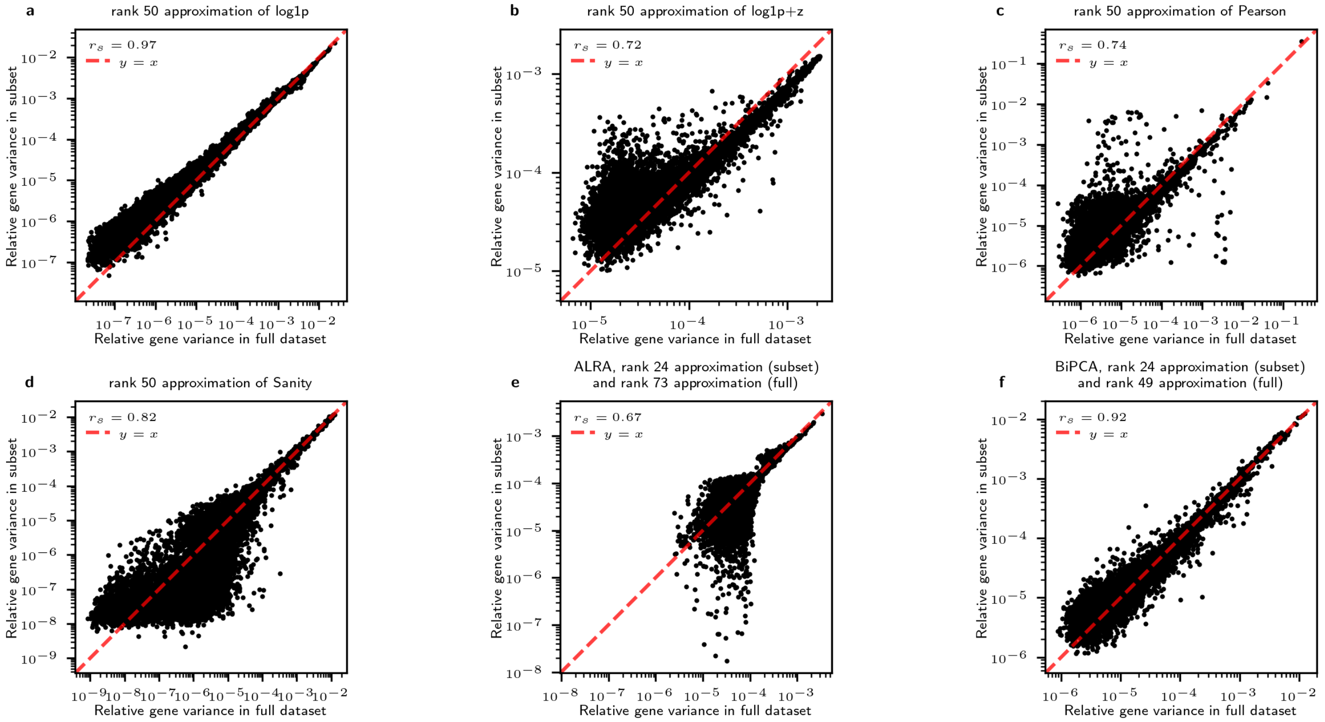


**Figure S10:** Comparison of the relative gene variances between the full dataset (x-axis) and the downsampled dataset (y-axis) for methods after low rank approximation (**a-f**). ALRA estimated rank 73 for the full dataset and rank 24 for the downsampled data (**e**), and BiPCA estimated rank 49 for the full dataset and rank 24 for the downsampled data (**f**).
